# A common molecular basis to the convergent evolution of the selfing syndrome in *Capsella*

**DOI:** 10.1101/653139

**Authors:** Natalia Joanna Woźniak, Christian Kappel, Cindy Marona, Lothar Altschmied, Barbara Neuffer, Adrien Sicard

## Abstract

Whether, and to what extent, phenotypic evolution follows predictable genetic paths, remains an important question in evolutionary biology. Convergent evolution of similar characters provides a unique opportunity to address this question. The transition to selfing and the associated changes in flower morphology are among the most prominent examples of repeated evolution in plants. Yet, to date no studies have directly compared the extent of similarities between convergent adaptations to selfing. In this study, we take advantage of the independent transitions to self-fertilization in the genus *Capsella* to test the existence of genetic and developmental constraints imposed on flower evolution in the context of the selfing syndrome. While *C. rubella* and *C. orientalis* have emerged independently, both have evolved almost identical flower characters. Not only the evolutionary outcome is identical but, in both cases, the same developmental strategies underlie the convergent reduction of flower size. This has been associated with convergent evolution of gene-expression changes. The transcriptomic changes common to both selfing lineages are enriched in genes with low-network connectivity and with organ-specific expression patterns. Comparative genetic mapping also indicates that, at least in the case of petal size evolution, these similarities are largely caused by mutations at the same loci. Together, these results suggest that the limited availability of low-pleiotropy paths predetermine closely related species to similar evolutionary outcomes.

## Introduction

Phenotypic changes evolved in response to changes in selective pressures or as a result of genetic drift. The influence of stochastic, deterministic and contingent factors in defining the route of phenotypic evolution is still debated^1^. Yet, in nature, there are many examples in which different populations have evolved identical features when confronted with similar ecological challenges^2,3^. These similarities are especially intriguing when they do not merely reflect the phylogenetic relatedness of individuals. One explanation that has been proposed is that these features represent optimal, or maybe even the only possible adaptations to particular environmental constraints^4^. This does not, however, appear to be a universal feature since many species have also adapted to the same environment through different mechanisms^1^. Other factors, such as the genetic and developmental constitutions of species may therefore also influence the repeatability of phenotypic evolution. In agreement with a role of evolutionary history in predisposing adaptive paths, meta-analyses of described events of repeated evolution have indeed suggested that convergent morphological adaptations seem to be more frequent in closely related lineages^5^. Because species that share ancestry have accumulated the same mutations and evolved the same developmental programs on which future adaptations will be built, they may be expected to respond similarly to new selective pressures. Additionally, the existence of shared standing variation in potentially adaptive traits may also underlie the repeated evolution of similar phenotypes^6^. Yet, the extent to which these different factors explain the convergent adaptations of closely related species, is not well understood.

Phenotypes are controlled by a series of genes that regulate each other through transcription or post-transcriptional interactions^7^. The gene regulatory networks (GRNs) controlling different traits are themselves interconnected and these connections will be modified during ontogenesis to ensure the successful orchestration of developmental programs. The complex structure of these networks may itself limit the genetic solutions to adaptation and predetermine evolutionary outcomes^8,9^. For instance, highly connected ‘hub’ genes play an essential role in network connectivity and, thus, any perturbations in their activities can have drastic pleiotropic consequences and strongly impair the fitness of organisms^10–12^. For this reason, only variations in a few genes may provide enough adaptive advantage in the face of new conditions because they allow specific changes in phenotypes and limit the pleiotropic consequences^13^. If such genes are limited, they are likely to constitute evolutionary hotspots, explaining the similarities observed in independent convergent evolution. In support of this prediction, a number of studies have now succeeded in identifying genes repeatedly involved in independent evolution of the same phenotype^13–17^. What these studies have also indicated is that the connectivity within GRNs is changing during ontogenesis and thus the target of selection within a given GRN is highly dependent on the developmental context^18^. Also, contrary to the above prediction, the perturbation of highly connected regulatory hubs has been shown to provide rapid evolutionary advantages by promoting phenotypic diversity^19^. Furthermore, not all networks appear to have a limited solution to evolve. In freshwater stickleback, for instance, pharyngeal tooth number has evolved independently through different means^20^. Identifying GRNs involved in convergent evolution may, therefore, provide important insights on how network structure can constrain phenotype evolvability. In particular, the hotspot genes identified by now have mostly come from studies of genetically simple traits with limited numbers of genes involved, such as flower colour or animal pigmentation ^13^. By contrast, much less is known about the genetic basis of convergent evolution of highly polygenic traits, with organ size being one prominent example.

The transition from outcrossing to selfing in plants offers a unique opportunity to investigate the genetic bases of convergent evolution of polygenic aspects of morphology. Self-fertilization is believed to be selected when compatible mates are rare and, thus, when its benefits in terms of reproductive assurance, outweigh the cost of inbreeding depression^21,22^. This transition has occurred independently hundreds of times during plant evolution and, in many cases of animal-pollinated species, it has been followed by very similar changes in flower morphology and function^23^. These similarities are such that these changes have been termed the selfing syndrome^24^. They include, in the predominantly selfing lineages, a strong reduction of flower size, a reduced pollen to ovule ratio and a decrease in the production of nectar and scent. The frequencies of these events and the resulting similarities in terms of phenotypic evolution allow challenging the above hypotheses regarding the influence of phylogenetic distances on the repeatability of evolutionary paths. Yet, no studies have directly compared the extent of similarities in the independent evolution of the selfing syndrome.

The genus *Capsella* has emerged as an ideal model to study the phenotypic consequences of the transition toward self-fertilization^25^. In this genus, two independent transitions to selfing have occurred both presumably through the breakdown of the selfi-ncompatibility system^26^. In a western lineage, the predominantly self-fertilizing *C. rubella* evolved from the outbreeding ancestor *C. grandiflora* within the last 200,000 years^27–29^. An earlier and independent event in an eastern lineage gave rise to *C. orientalis* from a presumed *C. grandiflora-like* ancestor within the last 2 million years^26,30^. The genetic basis of selfing syndrome evolution has been extensively studied in *C.rubella*^25,31^. In this species, a complex genetic basis underlies the evolution of flower size and pollen to ovule ratio. At least six mutations have contributed to reducing the size of the petals. Yet, most of the world-wide distributed accessions appear to share these mutations suggesting that the evolution of such traits has occurred early during the history of *C. rubella*, or at least before its geographical spread^25,31^. So far, nothing is known about the genetic basis of selfing syndrome evolution in *C. orientalis* and more particularly, on whether or not these two instances of convergent evolution rely on similar molecular mechanisms.

In this study, we compared the developmental, transcriptomic and genetic bases of the selfing syndrome in *C. orientalis* and *C. rubella.* In particular, we asked whether due to their close relatedness the phenotypic changes in these two species resulted from the same developmental and molecular mechanisms. We identified common transcriptomic changes between these independent instances of selfing-syndrome evolution and studied the pleiotropic properties to determine if their repeated involvement may have been predicted from network structures.

## Results

### Convergent evolution of flower morphology after the transition to selfing in the *Capsella* genus

To determine the extent of similarities in the phenotypic changes that have occurred in both selfing lineages, we compared the flower morphology in *C. grandiflora* (*Cg*), *C. rubella (Cr)* and *C. orientalis* (Co). The size of different flower organs, as well as of vegetative organs, was quantified in five representative accessions for each species (**Table S1**). Petal area is about five times smaller in both selfers compared to the outcrosser (**Fig. 1B**). This reduction is consistently observed between the different accessions of the three species analysed (**Fig. S1**). Average leaf size is only slightly reduced in *Cr* and *Co* (12 and 20 % smaller, respectively). Furthermore, the leaf area is highly variable within each species and the leaves from some of the selfing accessions are as large as those of *Cg* (**Fig. S1**). Within the flower, organ size reduction is not limited to petals, but also affects other organs (**Fig. 1 and S1**). Sepals are 15 and 21% shorter in *Cr* and *Co*, respectively, while the petals are for both selfers two times shorter than *Cg*’s; and anther and carpel length are 25 % smaller in the selfers compared to the outcrosser. In both *Cr* and *Co*, the evolution of self-fertilization has therefore been accompanied by a reduction of floral organ size and in particular, a strong decrease in petal dimensions. Strikingly, the size of all flower organs is not significantly different between the two selfers.

**Fig. 1.**
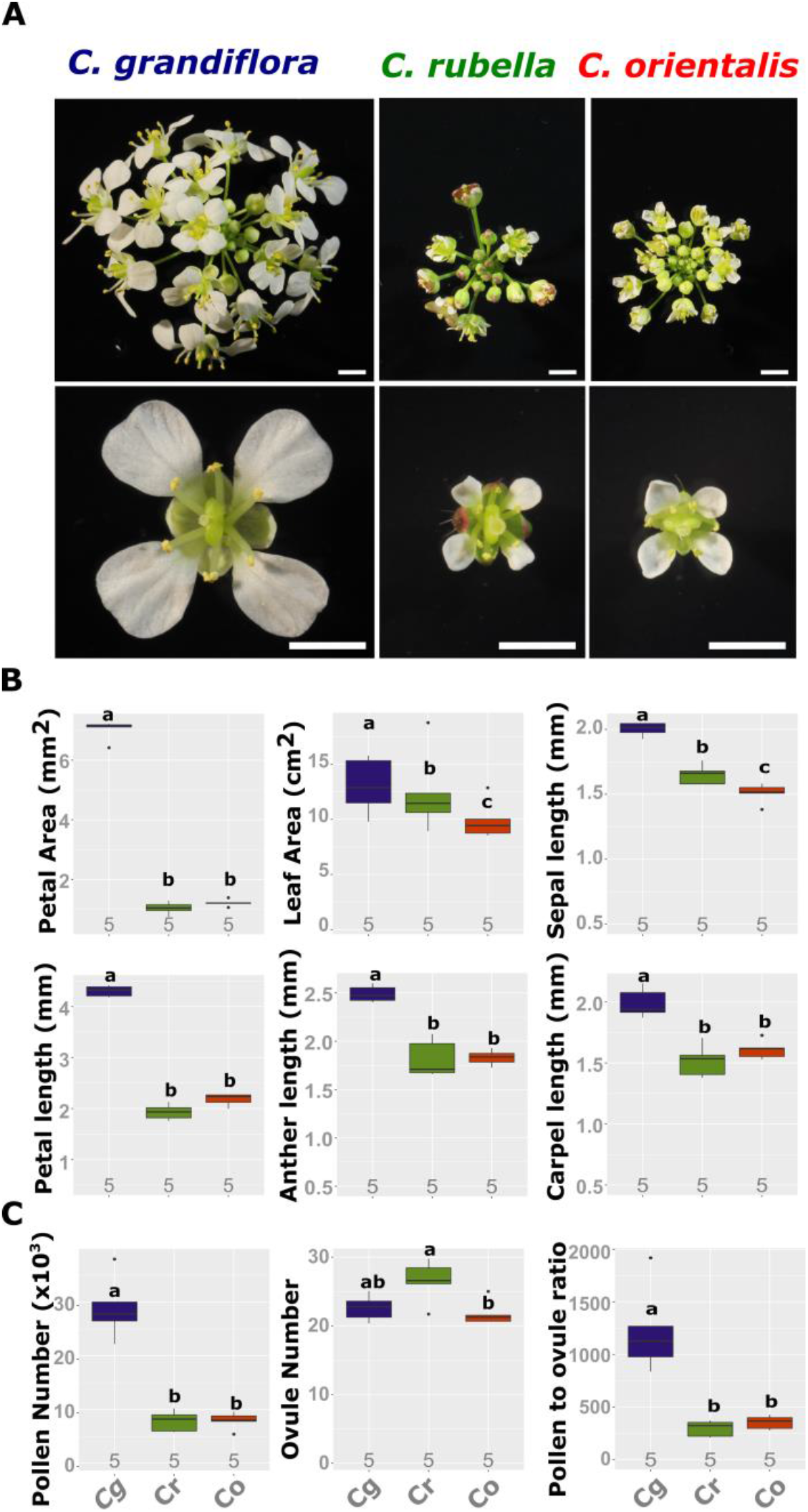
Convergent evolution of the flower morphology after the transition to selfing in the genus *Capsella*. (A) Photographs of an inflorescence and individual flower of *C.grandflora (*Cg*), C.rubella (Cr)* and *C.orientalis (Co)* are shown at the same mangnification. The scale bars indicate 3 mm. (B) Quantification of leaf size and flower organ size. (C) Quantification of sexual allocation. Pollen and ovule numbers were measured and used to calculate the Pollen to ovule (P/O) ratio. Letters indicate significant difference as determined by a Tukey’s HSD test. The numbers under each boxplot represent the number of independent accessions used for measurement (mean over individuals’ means from each accession was taken).

In many cases the transition to selfing is associated with a shift in the allocation of sexual resources. We therefore compared the number of pollen grains and ovules in the three *Capsella* species^24^. The pollen to ovule (P/O) ratio is reduced to a similar extent in both *Co* and *Cr* (**Fig. 1C**). This reduction is explained by a similar decrease, about 4 fold, in the number of pollen grains formed by each flower. As previously reported, we observe a slight increase, ~1.2 fold, in ovule number in *Cr*^25^. This increase is, however, not observed in *Co*, which develops a similar number of ovules as *Cg*. The increase in ovule number is relatively weak compared to the decrease in pollen number and has therefore a limited influence on the P/O ratio. As a result, the latter does not differ between both selfers.

Overall, these results indicate that despite their independent evolution *Co* and *Cr* have evolved very similar flower phenotypes. Furthermore, based on the comparison of leaf area between the three species, the changes in organ size seems to be largely restricted to flowers and most prominently to petals, indicating that organ-specific mechanisms are likely underlying these evolutionary changes.

### The same developmental mechanism underlies the reduction of petal size in *Cr* and *Co*

To determine whether similar development processes were responsible for the reduction of petal size in *Cr* and *Co*, we compared petal growth in each *Capsella* species using representative accessions (**Fig. 2 and S2**).

**Fig. 2.**
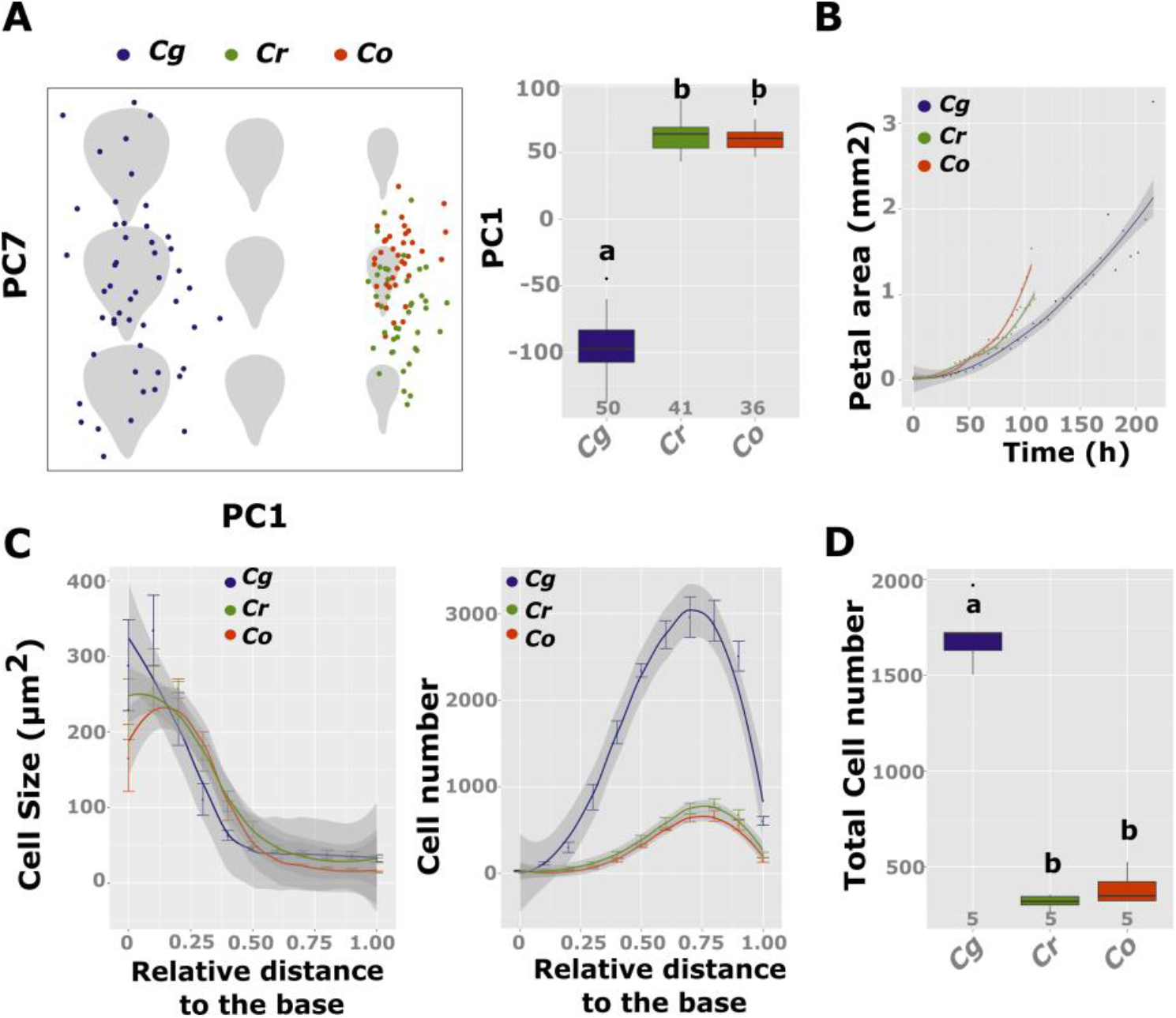
Similar developmental mechanisms underlie petal size reduction in both *C. rubella* and *C. orientalis.* (A) Principal component analysis on Elliptic Fourier Descriptors of the petal outlines. The PC1-PC7 morphospace is shown on the left where each dot represents the value of a single petal. The comparison of the PC1 values between each species is shown in a box plot on the right. The letters indicate significant difference as determined by a Tukey’s HSD test. The number of samples used is indicated under each boxplot. (B) Developmental series of petal growth in *C. grandiflora, C. rubella* and *C. orientalis* (n =4). (C) Average cell size (left) and cell number (right) along the petal longitudinal axis for the three *Capsella* species. The values are means of five replicates ± sem. Fitted curves are displayed and the dark-grey area around each line represents the 95% pointwise confidence interval. (D) Total cell number in the three *Capsella* species. Letters indicate significance as determined by a Tukey’s HSD test. The numbers of samples are indicated under each boxplot.

Because the final organ geometry reflects the sum of different growth patterns that occurred during development, it would be expected that if petals have been reduced through different mechanisms in the two selfer lineages, their overall shape should also differ. We therefore compared the petal geometry of the different *Capsella* species using principal component analysis (PCA) on Elliptic Fourier descriptors (EDFs) of the petal outlines. This analysis indicated that 74.3 % of the total variance could be explained by a single principal component, PC1 (**Fig. 2A**). This PC reflects a proportional variation in the overall petal size with a stronger modification along the transversal axis (**Fig. 2A**). PC1 clearly discriminates *Cg* from *Cr* and Co (Kruskal-Wallis test, p= 2.2 x10^-16^, **Fig. 2 A**), but fails to separate *Co* from *Cr.* Only PC7 and PC8, which each only explained 0.1% of the total variance, are significantly different between *Cr* and Co. However, these two PCs reflect only minor shape variation of the outline curvature at the junction between the petal limb and claw, and are not significantly different between each of the selfers and the outcrosser (**Fig. 2**). The difference in PC7 and 8 between *Cr* and *Co* is, therefore, more likely to be due to phenotypic variation in the ancestral outcrossing population rather than independent adaptation to self-fertilization. Overall, these results indicate that the two selfers have evolved similar petal size and shape suggesting analogous underlying developmental processes.

In plants, the final organ size is mostly determined by two processes, cell proliferation and cell elongation. Any variations in the rate or duration of these processes will therefore influence the final organ dimension. Following the growth of petal primordia over time in a representative accession for each of the species did not reveal any difference in the initial petal growth between *Cg*, *Cr* and *Co* (**Fig. 2B**). However, the petals cease to grower earlier in *Cr* and *Co* compared to *Cg* indicating that the duration rather than the rate of growth has been modified in the selfers. No difference in average cell size per segment along the longitudinal axis of the petal was observed between the three species (**Fig. 2C**). The numbers of cells in each section, however, strongly differ between the selfers and the outcrosser (**Fig. 2 C**). *Cg* petals consist of about 4.5 times more cells than those of *Cr* and *Co* (**Fig. 2D**). The total cell number of both selfers is not significantly different and the number of cells follows an identical pattern along the longitudinal axis. The two selfers have therefore evolved petals constituted by the same number of cells.

Despite the independent evolution of *Cr* and *Co* lineages, the same developmental mechanism underlies the reduction of the petal size, i.e. a reduced cell number due to shortening of the cell proliferation period.

### Convergent evolution of gene expression in *Cr* and *Co*

In many cases, the morphology of organisms evolved as a result of changes in the expression and/or mRNA abundance of genes regulating developmental programs^14,32,33^. The reduction of flower size in *C. rubella* is no exception and the underlying polymorphisms identified so far affect the mRNA quantity of key growth regulators^34,35^. Given the developmental similarities between *Cr* and *Co*, we hypothesized that their morphological adaptations to selfing may have been caused by mutations affecting the same gene regulatory networks (GRNs). If this was the case, it would be expected that the convergent morphological evolution in these lineages has also been accompanied by similar changes in gene expression.

To test this hypothesis, high-throughput RNA sequencing (RNA-seq) was performed in different tissues of the three diploid *Capsella* species. These tissues included 10-day old seedlings, young flower buds undergoing intense cell proliferation (later called young-flower) and old expanding and maturing flower buds (hereafter named old-flower). PCA using vst normalized read counts as well as hierarchical clustering based on Euclidean distances showed a clear grouping according to tissue type, developmental stage and evolutionary distance. PCA also allowed to identify differences based on mating system (**Fig. 3A** and **S3A**). PC1, which explained 21% of the total variance, separates the samples according to their tissue of origins. Flower samples clustered together at low PC1 values, while seedling samples show higher scores. PC2 discriminates the samples according to their developmental stages, with actively dividing tissues, such as young-flowers, having higher PC2 values than old-flowers and seedlings, which have a lower proportion of dividing cells. PC3 separates the samples according to their phylogenetic relationships, with *Co* being further apart from both *Cr* and *Cg.* PC4, however, distinguishes the samples according to their mating system with *Cr* and *Co* having lower PC4 values than *Cg.* This separation suggests that similar transcriptomic changes have occurred in both selfing lineages. To better quantify the extent of overlap between the transcriptomes of the two selfers, we performed a differential gene expression analysis comparing first each selfer to the outcrosser before overlapping the lists of Differentially Expressed Genes (DEGs). Out of 26521 genes, we found a total of 8242 (3097 upregulated and 5613 downregulated genes in at least one sample) and 10281 (4047 upregulated and 7021 downregulated in at least one sample) DEGs in *Cr* and *Co*, respectively (**Fig. 3B** and **S3**). Among them, 6205 genes were called differentially expressed in both selfers, of which 4173 (~ 70%) genes show changes in the same direction relative to the outcrosser in at least one tissue. We later refer to these genes as coDEGs. To determine whether this convergent evolution was also associated with correlated changes in gene expression levels, we compared the fold changes in transcript abundance of the coDEGs in *Cr* and *Co* (**Fig. 3C**). Gene expression changes were globally and significantly correlated (Spearman’s r = 0.76, p-value < 2.2e-16) suggesting that the expression of the same genes has also evolved to a similar magnitude in both selfing lineages. Consistent with the phenotypic changes that occurred in selfers, coDEGs were enriched for many different Gene Ontology (GO) terms including developmental processes involved in reproduction or metabolic processes (**Fig. S4**). The repeated evolution of flower morphology has therefore been accompanied by convergent evolution of gene expression.

**Fig.3.**
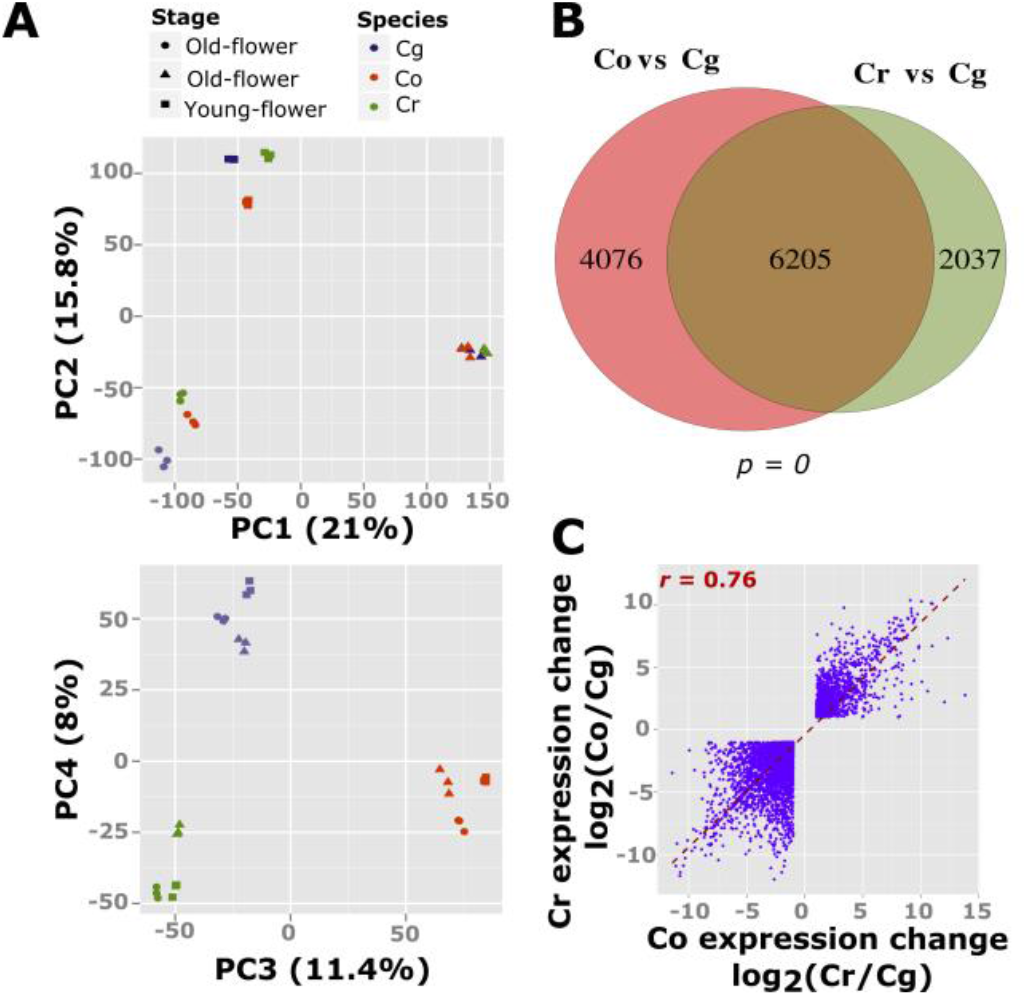
Convergent evolution of gene expression after the transition to selfing in the genus *Capsella.* (A) Principal component analysis factorial map illustrating the distribution of the species and tissues transcriptomes along the largest components of variance. Each symbol represents a biological replicate. The symbol shape and colour legend is indicated on the top. The percentage of variance explained by each component is indicated on the axis titles. (B) The genes differentially expressed between *Cr* and *Cg* are compared to those differentially expressed between *Co* and *Cg*. The significance of the overlap was calculated using a hypergeometric test. The *p*-value is indicated under the Venn diagram. (C) Correlation in gene expression changes between the *Cr* and *Co* DEGs. The Spearman correlation coefficient *r* is indicated on the graph.

### Increased convergent gene-expression evolution in flowers

Because the phenotypic changes in selfers are mostly apparent in reproductive organs, we tested whether the convergent evolution of gene expression was restricted to the flower transcriptomes. DEGs were identified in all the tissues analysed (**Fig. 4A**). We found a total of 6648 DEGs, in at least one selfer compared to the outcrosser, in the seedling samples, 7854 in the young-flower samples and 8507 in the old-flower samples. A significant number of genes were called differentially expressed in both selfers in all samples and rather few of them were found to be sample-specific (**Fig. 4A** and **S5A**). Nevertheless, interspecific comparisons based on pairwise gene-expression differences revealed a longer gene-expression branch length between the selfers and the outcrosser in the flower samples compared with the seedling samples (**Fig. 4B**). In both *Cr* and *Co,* the young-flower and old-flower samples show higher gene-expression changes compared to seedlings and the overlaps between the DEGs in *Cr* and *Co* were also higher in the floral tissues (**Fig. 4A**). Consistently, when focusing our analysis on genes changing in the same directions in both selfers, we observed that about 70% of the coDEGs were co-differentially expressed in the reproductive tissues. Selecting genes differentially expressed in only reproductive samples, only in vegetative tissues or in both, revealed that 76% of the coDEGs correspond to organ-specific changes, 90% of which are specific to flowers (**Fig. 4A** and **C**). Only 15% were co-differentially expressed in all samples and the remaining 9% of coDEGs are differentially expressed in both organ types in only one of the selfers (not displayed on the donut plot; **Fig. 4C**). The fold change in gene expression level was, however, similarly correlated for the coDEGs in both vegetative and reproductive tissues (Spearman’s r = 0.69 and 0.68, respectively). Overall, these results indicate that most of the convergent gene-expression evolution has occurred within the flowers and suggest an accelerated evolution of the flower transcriptome in selfers.

**Fig.4.**
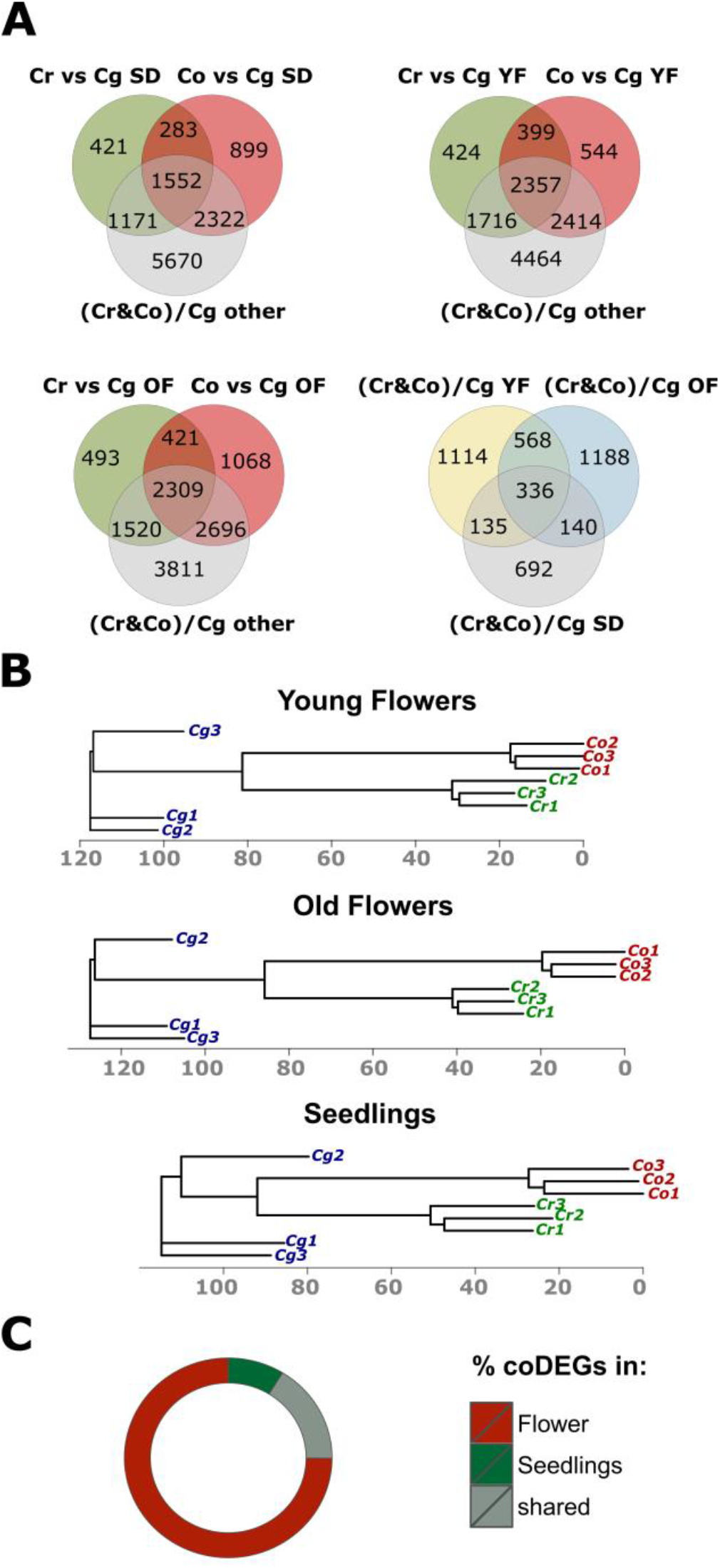
Higher divergence In the flower transcrlptome after the transition to selfing. (A) The genes differentially expressed between *Cr* and Cg are compared to those differentially expressed between *Co* and *Cg* for each tissue type. (SD: seedling; YF: young flowers; OF: old flower). On the bottom right, the list of genes called differentially expressed in both selfers ((Cr&Co)/Cg) are compared between tissue-type. (B) Neighbour-joining tree based on Euclidean distances calculated using vst normalized read counts. Note that the branch separating the selfers from *Cg* is longer when the species comparison uses the flower transcriptome. (C) Donut plot illustrating the proportion of genes differentially expressed in the same direction in both selfers within each tissue-type. Additional 9% of coDEGs differentially expressed in both organ types in only one of the selfers is not displayed on the plot.

### Low-pleiotropic organ-specific GRNs underlie the convergent evolution of the selfing syndrome

Organ-specific evolution, such as the reduction of flower size after the transition to selfing, can arise through changes in tissue-specific regulators or through tissues-specific modifications of general regulators. Gene ‘reuse’ in convergent phenotypic evolution has often been proposed to be explained by the fact that only a few genes may have a specialized function that allows ‘optimizing’ the phenotype of specific organs while avoiding pleiotropic effects^13^. We therefore sought to analyse the expression pattern and pleiotropy of the coDEGs. Their expression pattern across the samples was globally positively correlated between selfers and outcrossers (**Fig. 5A** and **S6A**). Only a small proportion of the genes appear to have a lower correlation coefficient suggestive of an evolution of their pattern of expression (**Fig. S6B**). We next asked whether coDEGs tend to have an organ-specific expression pattern. To this end, we determined the proportion of genes expressed in an organ-specific manner, which we defined as the genes whose expression is enriched by at least 5-fold in a specific tissue, in the coDEGs and the non-coDEGs (genes called differentially expressed in only one selfers, also called: Others) (**Fig. 5B**). The coDEGs are enriched in organ-specific genes (63%) compared to the non-coDEGs (31%). Consistent with the observation that most of the coDEGs are co-differentially expressed in flowers, 47% of the coDEGs show flower-specific expression whereas only 15% are preferentially expressed in seedlings. This pattern was further reinforced when the analysis focused on genes co-differentially expressed only in flowers. Indeed, within the flower coDEGs 56% were mostly expressed in flowers. In contrast, in the seedlings only 23% coDEGs correspond to genes mostly expressed in reproductive tissues, while 31% were preferentially expressed in seedling (against 15% in the flower DEGs). This data are therefore more consistent with the recurrent evolution of organ-specific GRNs rather than repeated changes in gene expression patterns.

**Fig.5.**
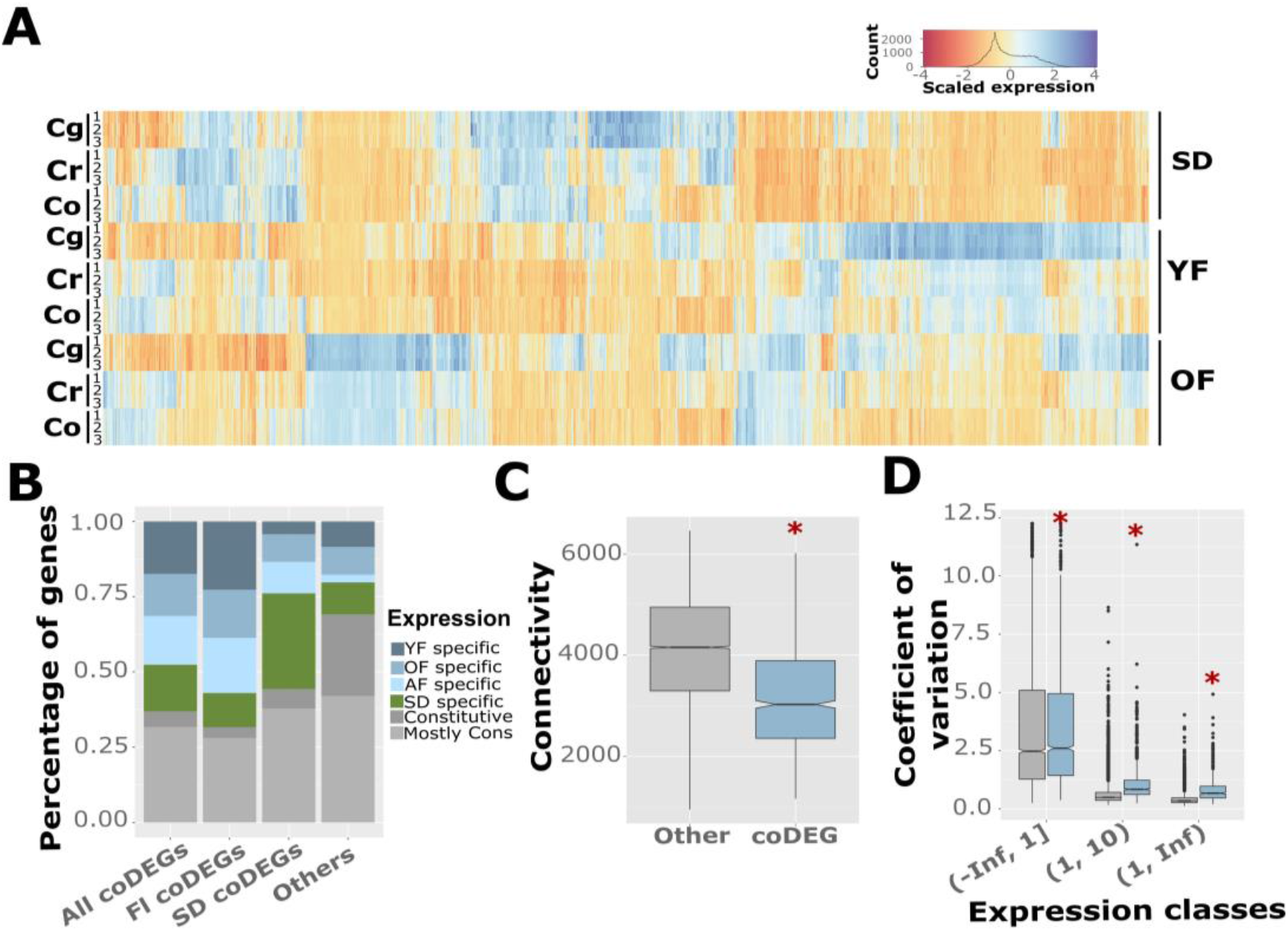
Low-pleiotropic gene regulatory networks underlie the convergent evolution of flower morphology. (A) Heatmap illustrating scaled expression values for differentially expressed genes going in the same direction in *C. rubella* and *C. orientalis* at one or more developmental stages. (B) Percentage of genes expressed preferentially at one developmental stage within the genes differentially expressed in both selfers compared to non coDEGs. YF indicates the proportion of genes, which are specifically expressed in Young flower, OF in old-flowers, AF in all flower samples, SD in seedlings. The proportion of genes with equal expression in all samples is shown in dark grey (‘constitutive’) and those with only weak differences in expression (between 1 and 5 fold) are shown in light grey (‘mostly cons’). (C) Boxplots showing the sums of network connectivity within a Cg population (as determined by^36^, 12896 quantified genes) for coDEGs and all other genes. Asterisks indicate significance (p < 0.05) determined by a Wilcoxon test. (D) Boxplots showing coefficients of variation of gene expression levels by expression group within the same Cg population. Expression categories for coDEGs (blue) and all others genes (grey) are based on averages over all samples, (Inf,1] corresponds to genes with tags per million (TPM) up to 1, (1,10] with TPM between 1 and 10, (10,Inf] indicates genes with TPM above 10. Categories roughly contain the same number of genes. Asterisks indicate significance (p < 0.05) determined by a Wilcoxon test.

The strength of connections between genes, also termed ‘network connectivity’, can be used to estimate gene pleiotropy from transcriptome data sets, as a gene having higher connectivity is more likely to have a pleitropic effect^36^. We therefore used the measures of connectivity of Josephs et al, (2017)^36^ extracted from the genome-wide analysis of gene expression in about 150 *Cg* individuals to test whether coDEGs tend to have low pleiotropy. coDEGs show a significant reduction in the sum of connectivity when compared to the non-coDEGs (**Fig. 5C**). This pattern was, however, not observed when randomly selecting 2000 genes, suggesting that the list of coDEGs are particularly enriched in genes with low pleiotropy (**Fig. S6C**). Consistently with previous results indicating a negative association between network connectivity and non-synonymous divergence, the coDEGs show a significant increase in dN/dS ratio that was not observed for the list of 2000 randomly drawn genes (**Fig. S6E** and **S6F**^36^). The coefficient of variation calculated from genome-wide expression data of *Cg* is also significantly higher for the coDEGs especially for genes with high expression value (**Fig.5D, S6D** and **S6G**). No change in coefficient of variation was observed when randomly drawing 2000 genes. Thus, these observations suggest that the two independent transitions to selfing have ‘reused’ genes under weak functional constraints, with low pleiotropy and variable expression in the ancestral population.

### A common genetic basis to the independent reduction of petal size in *Capsella*

The similarities in the phenotypic and molecular evolution of the two selfing lineages raise the question of whether they may be caused by recurrent mutations in the same genes. To test this hypothesis, we generated *Co* x *Cg* and *Co* x *Cr* interspecific hybrids using ovule rescue. From the F1 populations, we selected a single individual for each cross type and harvested F2 seeds. We therefore first sought to analyse the segregation of selfing syndrome traits in these populations to test for a common genetic basis. Indeed, if the same genes underlie the independent evolution of the selfing syndrome, the two selfers would carry mutations at the same genomic positions and, thus, selfing syndrome traits should not segregate in the progenies of *Co* x *Cr* F1 hybrids. In such case, the phenotypic distributions of such population should overlap with those of the two parents. If, however, the change in flower morphology was caused by mutations in different loci, their phenotypic distributions in the same F2 populations would be expected to transgress beyond the parental values. We therefore analysed the segregation of petal size as well as ovule and pollen numbers in the *Co* x *Cr* F2 populations (**Fig. 6** and **S7**). Both of the selfers show very similar narrow distributions of petal area characterized by coefficients of variation (CV) of 0. 125 and 0.19, for *Co* and *Cr* respectively. The petal size distribution in the outcrosser was characterized by a similar CV of 0.2 but with a ~5 times higher mean. The F2 progenies of crosses between either of the selfers with the outcrossers produced a much wider distribution (CV = 0.34 and 0. 27 for *Cr* x *Cg* and *Co* x *Cg* respectively) which covers the entire range between the parental strains. In contrast, the petal size distribution was narrower (CV= 0.2) in *Co* x *Cr* F2 hybrids with a mean centered between those of the two selfer parents (**Fig. S7**). We detected transgressive segregation beyond the higher parental values but not on the lower side of the distribution. More importantly, no large flower phenotypes were observed in the *Co* x *Cr* F2 population. We next compared the frequency distribution of petal size observed in this F2 population with frequency distributions simulated with different QTL models having an increasing number of contributing loci (**Fig. S8**). The effect and location of the loci used in the simulations were defined based on previous QTL studies and the distribution was centred around the mean between the two selfers using previously estimated residual variance components^25,37^. The observed phenotypic distribution was not statistically different from those simulated with the models using one or no segregating petal size QTL. The two distributions became different only when the model uses at least three segregating loci. From this point, the spread of the distribution was higher in the simulated data compared to the observed one (CV = 0.3 for simulated distributions from models with 3 QTL). This is consistent with only a few loci influencing petal size in the *Co* x *Cr* F2 population and suggests that part of the genetic basis underlying the reduction of petal size is shared between the two selfers. The frequency distribution of ovule and pollen numbers in *Co* x *Cr* F2 population was, however, much wider (**Fig. S7**). We found significant transgressive segregation for both of these traits. The segregation of ovule numbers was best explained by a model with at least 12 segregating QTL and the pollen number distribution fits more accurately the distribution from models with at least 16 segregating QTL. These results suggest therefore that a large number of loci influencing these traits are segregating in the *Co* x *Cr* F2 population suggesting a different genetic basis in both selfers.

**Fig. 6.**
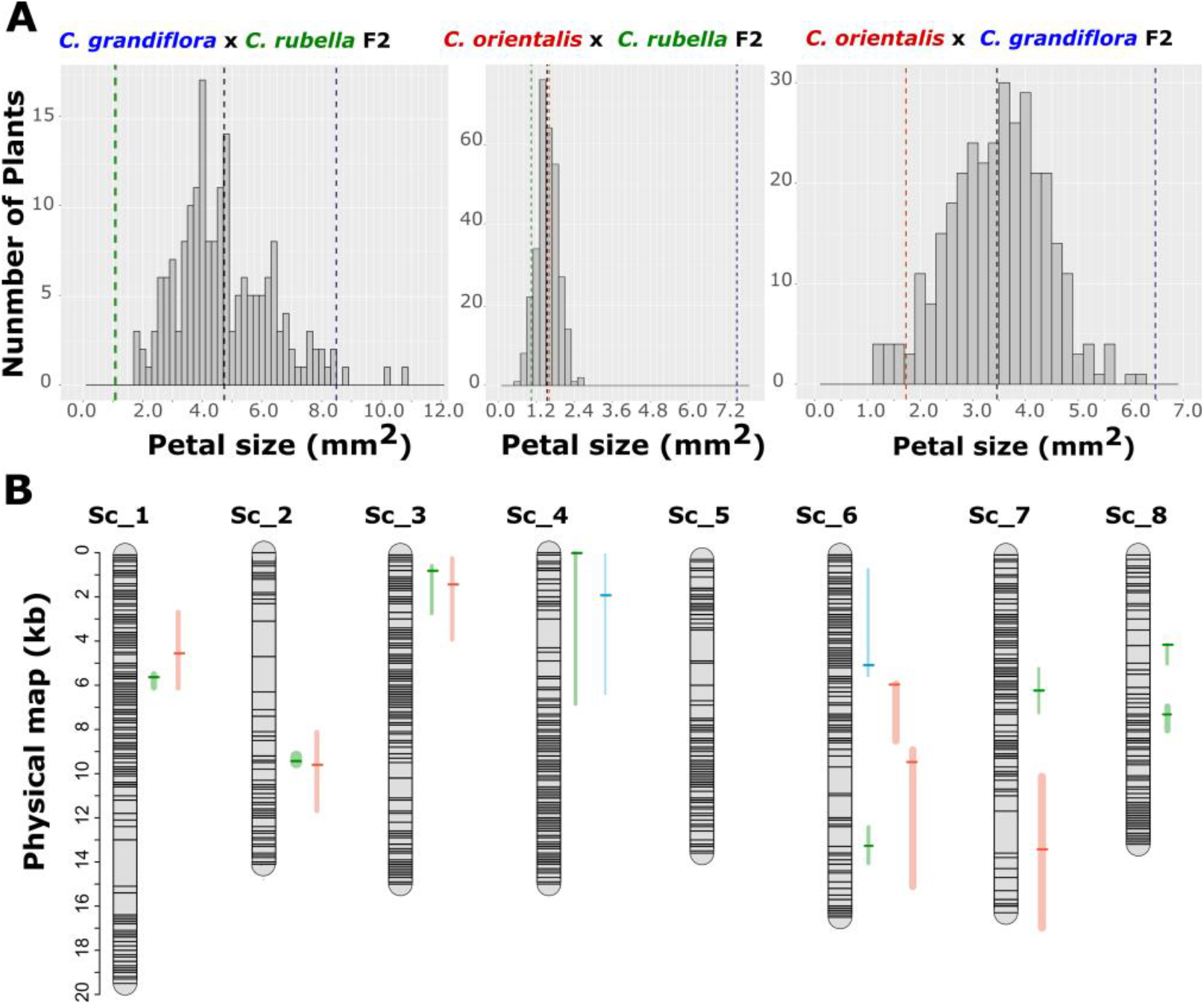
Genetic basis of the reduction of petal size after the transition to selfing in *Capsella.* (A) Petal area frequency distribution in *Cg* x *Cr*, *Co* x *Cr* and *Co* x *Cg* F2 population. (B) Comparison of the genetic basis underlying the reduction of petal size in *Cr* and *Co*. The positions of QTL influencing petal size are indicated on the physical map. The QTL identified in this study in the *Co* x *Cg* population are indicated in green, in the *Co* x *Cr* are indicated in blue. Those previously identified in the *Cr* x *Cg* RILs population^25^ are shown in red. The LOD scores peak are indicated by horizontal lines and the vertical lines represent the 1.5 LOD score confident interval. The width of the vertical lines illustrates the strength of the QTL.

To confirm these observations, we used the population generated above to map the loci involved in the reduction of petal size in *Co*. A total of 462 F2 individuals of the *Co* x *Cg* and 381 F2 individuals of the *Co* x *Cr* populations were genotyped using double-digest restriction site associated DNA (ddRAD) sequencing and phenotyped for vegetative and reproductive traits. The genotypes obtained were then used to establish genetic maps using the Kosambi mapping function. As expected from previous studies, this resulted for both populations in eight linkage groups which were in good agreement with the position of the markers on the genome sequence^38^ (**Fig. S9** and **S10**). ddRAD markers were globally evenly distributed throughout the genome and only weak segregation distortion was observed along the 8 linkage groups. We therefore used this genotype information to map QTL influencing the measured traits (**Fig. S9** and **S10**). A multiple QTL approach was used to identify the most informative QTL models, which were then used to position the QTL within the genome (**Table S2** and **S3**). The positions of the QTL identified were then compared to previous QTL mapping experiments in *Cr* x *Cg* F2 and RIL populations (**Fig. 6** and **S11**^25,37^). Only one QTL, SIQTL1, influencing the self-compatibility in *C. orientalis* was identified. This QTL overlaps with the *Capsella* S-locus which contains the S-locus reporter kinase and the linked S-locus Cys-rich proteins and which has been reported to underlie the loss of self-incompatibility in *C. rubella*^25,27,37^(**Fig. S11**). Consistently, self-incompatibility was not segregating in the *Co* x *Cr* F2 population. This further supports the observation that mutations at the same locus seem to underlie the loss of self-incompatibility in *Cr* and Co^30^. For petal area, we identified 8 QTL which together explained 53% of the total phenotypic variance *Co* x *Cg* in population (**Fig. 6**). Most of these QTL act additively as we only detected a significant interaction between PAQTL5 and PAQTL8 (**Table S2**). Notably, three of these QTLs, which together explain more that 30% of the total phenotypic variance, overlap with QTL previously identified in two other independent *Cr* x *Cg* populations^25,37^ (**Fig. 6** and **S11**). In particular, the QTL on scaffold two which here explained by itself 23% of the total phenotypic variance *Co* x *Cg* population was also found to have a major influence on *Cr* petal size. From the remaining QTL, only at PAQTL4, 6 and 7, which together explained 11 % of the total phenotypic variance, the *Cr* alleles reduced the petal size and are therefore likely to underlie lineage-specific modifications after the transition to selfing (**Table S3**). Consistent with the above segregation analysis, this QTL mapping experiment suggests that the loci with stronger effects may have contributed to the reduction of petal size in both of the selfing lineages. PAQTL 1 to 3 affected both the length and width of the petals, but none of them has an influence on leaf size, indicating that as in *Cr* these loci act in an organ-specific manner. As it could be expected, none of these QTLs were detected in the *Co* x *Cr* population (**Fig. 6** and **S11**).

We identified 5 and 3 QTLs influencing ovule number within the *Co* x *Cg* and *Co* x *Cr* F2 populations, respectively. Two of these QTLs, OQTL5 in *Co* x *Cg* and OQTL2 in *Co* x *Cr* were overlapping with the QTL identified in *Cr* x *Cg* populations (**Fig. S11**). The QTL models identified, however, explained only 20 % of the total phenotypic variance suggesting many other loci with a weak effect were also segregating in this population (**Table S2**). Similarly, no significant QTL influencing pollen number were detected in these populations suggesting, here as well, the contribution of a large number of small effect mutations. Such scenario is consistent with the above segregation analysis, which revealed that the observed phenotypic distribution could be explained by a complex genetic basis.

## Discussion

We sought to analyse the extent of similarities in the phenotypic evolution that have followed independent transitions to selfing in two closely related species. Remarkably, our results indicate that the two selfers have evolved almost identical flower size and shape. This has occurred through the same developmental mechanisms, a shortening of the duration of the cell proliferation period. As a result, these two species have evolved petals consisting of the same number of cells. The similarities were not only restricted to flower size, but both of these species also showed a similar decrease in the number of pollen grains. The number of ovules was, however, only increased in *Cr.* The level of similarities in the phenotypic changes of the two selfers is intriguing and raises the question of what are the underlying causes.

### Ecological value of selfing syndrome traits

These similarities could be explained by the fact that these specific trait values confer a strong phenotypic advantage in a selfing context. Strong reduction in both petal size and pollen numbers are prominent recurring features of the selfing syndrome^25^. Theoretical modelling and the rapid evolution of selfing syndrome observed in recently emerged selfer lineages support the idea that these phenotypic changes could evolve as a result of positive selection, rather than the relaxation of constraints imposed by the needs of attracting pollinators^27,28,39,40^. Flower size has been shown to affect the distance between anthers and stigma, also known as herkogamy^25,41^. Variation in herkogamy inversely correlates with the ability of plants to self-fertilize, most likely because it facilitates the deposition of self-pollen onto the stigma^42–46^. A reduced flower size may therefore act as mating system modifiers by facilitating the establishment of self-fertilisation^47,48^. Nevertheless, flower size has been shown to be reduced through different mechanisms and to a different extent in different selfing species^24^. However, here both selfers have evolved exactly the same petal morphology.

Moreover, they have done so through the same developmental strategy and our genetic and transcriptomic analyses suggest that they have, at least in part, used similar genetic paths. The large extent of similarities may therefore not only be driven by the adaptive value associated with petal size, but also because a limited number of genetic solutions exist to reduce flower size without having detrimental consequences on plant fitness.

Compared to flower size, it is less obvious how pollen number would act as a mating system modifier and contribute to selfing evolution. It is more likely that the reduction in pollen number reflects the re-allocation of the resources invested in male function in out-crossing species^47,48^. Here, our genetic analyses indicate a different genetic basis in the two selfing lineages with the contributions of several small effect mutations. This would be consistent with this trait evolving as a consequence rather than a cause of selfing evolution, in which case mutations with small effect refining the fitness optimum would be expected to contribute^47^. That said, why would these two species evolve the same number of pollen grains? Pollen to ovule ratio has been shown to correlate with mating systems and ecology^24,49,50^. It is therefore conceivable that an optimal ratio between pollen grains and ovules exists and that because *Cr* and *Co* share the same ancestry and life history characters their phenotype would converge towards the same values.

### The genetic constraints imposed on flower size evolution

Our study indicates that, to a large extent, the two selfing lineages share the molecular mechanisms underlying the reduction of petal size and, possibly, even rely on mutations within the same genes. This could be expected if, for instance, the adaptation to selfing in one species would have been helped by the introgression of ‘selfing’ alleles from the oldest selfing lineage. This seems, however, unlikely for several reasons. First, the lineage in which *C. orientalis* has evolved diverged from the *C. grandiflora/C. rubella* lineage about 1 to 2 MYA^26,51^. Furthermore, the species occupy nonoverlapping geographical ranges, which prevent any gene flow between modern populations^26^. Finally, no signs of recent introgression were identified at close proximity of shared polymorphisms between the two *Capsella* selfing lineages^29^. The similarities in these two instances of selfing-syndrome evolution may therefore suggest that specific regulatory nodes in the gene network controlling flower size are more suitable to adapt plant phenotypes to selfing.

Although a large number of pleiotropic growth regulators have been identified in plants, it is still unclear how many of them can be modified to change the size of a specific organ. In *Capsella,* the transition to selfing has been associated with a strong reduction of flower dimensions without strong effects on the overall body size. Mutations affecting petal-specific regulatory elements have been shown to play an important role in the *C. rubella* selfing syndrome^34^. It is therefore plausible that only the genes with similar regulatory architecture could allow to specifically reduce the size of the flowers. Consistent with the recurrent involvement of mutations having an organ-specific effect, we have observed that the gene-expression changes common to the two selfers tend to correspond to genes mostly expressed in flowers. Yet, it is still unclear how many of the plant growth regulators could lead to organ-specific phenotypic evolution.

The capture of standing genetic variants from the ancestral outcrossing population has contributed to the evolution of the selfing syndrome in *C. rubella*^34^ Furthermore, long term balancing selection has been shown to maintain polymorphisms over several millions years in *C. grandiflora*^29^. The availability of genetic variants having an adaptive value for selfers in outcrosser populations could therefore predetermine the path to selfing syndrome evolution. Indeed, if for instance purifying selection on ‘small flower’ alleles was less efficient at specific loci, it could explain why such loci would underlie independent evolutionary events. Cases of convergent evolution due to shared standing variation have been described before^6^. We also observed that the expression the coDEGs tends to be more variable in the outcrossing population. The similarities in the independent evolution of the selfing syndrome may therefore originate from the GRNs under weak functional constraints that may provide the necessary variation for a rapid track to phenotypic evolution.

### Pleiotropy of the genes involved in independent evolution of the selfing syndrome

Genes with low pleiotropy, yet strong effects on phenotypes are expected to be more suitable for phenotypic adaptation^13^. The convergent evolution in gene expression in the two *Capsella* lineages appears to affect genes with low pleiotropy. These genes are on average less connected compared to other differentially expressed genes and most of them are expressed predominantly at a particular developmental stage or in a particular organ type. Furthermore, these genes also show higher amino acid divergence and among them, genes having a high expression level also show higher coefficients of variation in gene expression in *Cg*. These observations are consistent with previous findings indicating that genes with lower connectivity are under weaker functional constraints^36^. Altogether, these observations are in agreement with the idea that genes with lower pleiotropy are more likely to underlie repeated instances of phenotypic evolution. Although it may be difficult to decipher what is the cause-consequence relationship, these results also suggest that the recurrent involvement of the same genes is also determined by the selective pressures imposed by the network structure. By having a more variable expression, the genes with low connectivity could provide genetic variation upon which evolutionary processes could act. The genetic variation caused by the reduced negative selection on low pleiotropic genes could therefore by itself predetermine the paths of evolution.

In summary, our results are in line with the predictions that independent phenotypic changes in response to the same ecological challenge are likely to be similar between closely related species that share ancestry^1^. Nonetheless, our study not only revealed similar evolutionary outcomes but also that very similar paths have been followed by the two lineages in response to selfing evolution. Gene ‘reuse’ in convergent evolution of flower morphology is associated with reduced levels of gene pleiotropy. Determining whether the constraints imposed on flower size are caused by limited genetic solutions to the evolution of specific phenotypes or by the availability of genetic variants to feed evolutionary processes will require further studies. By identifying loci involved in independent selfing syndrome evolution, this work offers for the first time the means to answer these questions.

## Materials and methods

### Biological materials and growing conditions

The geographical origins of the different *Capsella rubella (Cr), C. orientalis (Co)* and *C. grandiflora* (*Cg*) accessions used in this study are detailed in Table S2.1. *Cr1504, Co1983* and *Cg926* were used for the analysis of petal development and the comparative transcriptomics experiment. Interspecific hybrids were generated by crossing *Co1983* with *Cr4.23* and *Co1983* with *Cg926* followed by ovule rescue as previously described^52,53^. F1 plants were allowed to self for each cross-type to generate the F2 populations. The phenotyping analysis in the F2 population was conducted using the progenies of one F1 plant per cross type. Note that *Cr4.23* was used instead of *Cr1504* in the genetic mapping experiment because genetic incompatibility between *Co1983* and *Cr1504* strongly impaired hybrid performance^53^. Nevertheless, because genetic diversity is relatively low within *Cr* and the evolution of selfing syndrome has occurred before its geographical spread, both *Cr4.23* and *Cr1504* shared the same mutations and molecular mechanisms underlying the evolution of selfing syndrome^25,31,35,53^.

All the plants were grown under long day conditions (16h light/8h dark) at 70% humidity, with a temperatures cycle of 22°C during the day and 16°C during the night and with a light level of 150 μmol m^-2^s^-1^.

### Morphological measurements

To compare the species-wide phenotypes of *C. rubella, C. orientalis* and *C. grandiflora* five accessions for each species were analysed as listed in Table S2.1. For all these accessions and the F2 interspecific populations, the following characters were measured.

Leaf size was quantified from the fully expanded 12^th^ leaves. Flowers organs were measured from the 15th and 16th fully opened flowers on the main inflorescence stem. Three petals, three sepals and three stamens were measured for each plant. Carpels were collected from the mature 17^th^ and 18^th^ flowers before fertilization. Dissected flower organs were flattened and scanned at a resolution of 3200 dpi while leaf images were digitalized at 300 dpi (HP ScanJet 4370). Area, length and width of the different organs were measured from these digitalized images using ImageJ (http://rsbweb.nih.gov/ij/).

The number of pollen grains and ovule produced by each flower were estimated from two flowers per individuals. To estimate the number of ovules, the carpels from the 15^th^ and 16^th^ flowers were incubated in a clearing solution (0,2g chloral hydrate; 0.02g glycerol; 0.05ml distilled water) overnight. They were then mounted in the same solution on microscope slides and the number of ovules in each carpel was counted using a light microscope Olympus BX 51. Pollen numbers were estimated from the flowers 19^th^ and 20^th^ just before anther opening. After desiccation at 37°C overnight, the pollen contained in the anthers was released in a 5% Tween-20 solution through sonication. The number of pollen grains per flower was then determined using a hemocytometer under a light microscope, Olympus BX 51.

The morphometric analysis of petal shape was performed using Elliptic Fourier Descriptors (EDF)^54^. First, the digital images were converted into binary images using ImageJ^55^. Coordinates of the petal outlines were extracted using the bwboundaries function in Matlab. Outlines were then Fourier transformed^56^ using the base of the petal as reference. Petal shapes were then compared by performing a principal component analysis (PCA) on EDF coefficients using the R function prcomp^57^. The effect of the PCs was illustrated by reconstituting the petal outlines through inverse elliptical Fourier transformations^56^ and using the maximum and minimum PC scores. PCs discriminating the *Capsella* species were identified using a Kruskal-Wallis test with the formula “principal component score ~ species”.

The kinetic analysis of petal growth was performed by manually dissecting two developing petals from each flower buds, starting from the oldest unopened flower and extending toward the youngest buds until the petals could not be dissected. Petal images were acquired using an Olympus BX51 microscope and areas were quantified as described above. Average petal area was then plotted against time based on the average plastochron of each genotype; itself estimated from the number of flowers produced over a 7-day period. To investigate the cellular basis of differences in organ size, dried-gel agarose prints^58^ of whole petals were generated from 5 plants for each species’ representative accession. Cell outlines were imaged under Differential Interference Contrast (DIC) on an Olympus BX51 microscope using an AxioCam ICc3 camera (Zeiss). The Python module scikit-image was used to process the resulting images. Cell-outlines were segmented by adaptive thresholding. Binarized cell borders were then dilated, skeletonized and curated by overlaying them with the initial images in GIMP (https://www.gimp.org/). Cell areas and centroid coordinates were extracted and used to determine the cell numbers and average area per sections along the petal longitudinal axis. The comparison of cellular patterns in each genotype was performed using R ^57^.

### Transcriptome analysis

Total RNA was extractedusing RNeasy Plant Mini Kit (Qiagen) from 10-days old seedlings, dividing flower buds and expanding/maturing flower buds. Buds were considered in an active cell division period when they were younger than stage 10 according to the *Arabidopsis thaliana* nomenclature of developmental stages^59^. Older flower buds were pooled and considered to correspond to the maturation/expanding phase. Three biological replicates for each development stages and for each species were used. TruSeq RNA libraries were generated according to the manufacturer’s instructions (Illumina) and sequenced using the Illumina HiSeq2000 instrument (1 × 50 cycles).

RNA-Seq data were processed using Trimmomatic^60^ to remove adapter sequences. Quality control was done using FastQC (http://www.bioinformatics.bbsrc.ac.uk/projects/fastqc). Reads for all samples were mapped against the *C. rubella* reference genome (Cru_183 from phytozome.org) using TopHat2^61^. Quantification for *C. rubella* predicted genes was done using HTSeq-count^62^. Data analyses and illustrations were done using R^57^. Differentially expressed genes between each selfer and the outcrosser at each developmental stage were obtained using DESeq2^63^. Genes were considered to be significantly changed when adjusted p-values were below 0.05 and log2FC above 1. Reads counts were vst normalized for clustering and PCA using DESeq2. PCA was done using the R prcomp function. Neighbour-joining trees were constructed with the R ape package^64^ using pairwise Euclidean distances based on vst normalized read-counts. Analyses with gene connectivity and divergence measurements were done using publicly available data sets^36^. Expression values to calculate the coefficients of variation in a Cg population were obtained using Kallisto^65^, rawdata were downloaded from NCBI SRA, project number PRJNA275635. The identification of the gene categories overrepresented in the list of differentially expressed genes was performed using Gene Ontology (GO) enrichment analysis^66–68^.

### Phenotypic segregation analysis

Transgressive segregation was estimated as previously described^69^. Chi-square tests were used to compare the number of individuals expected to exceed the means of the parent by at least 2SD (calculated based on the number of individuals above this threshold in the parental populations scaled to the size of the F2 population), with the observed number of individuals exceeding those thresholds in the F2 population, where SD was the pooled standard deviation of parental populations. Frequency distributions were simulated with various QTL models using the sim.cross function of the R/QTL package add-ons implemented in R ^57,70,71^. The effect and position of the QTL, as well as the genetic map used in the simulation, were based on previous QTL mapping experiments in *C. rubella*^25,31^. The sim.cross function was modified to centre the distribution around the mean between the two selfers average using previously estimated residual variance for each of the trait investigated^25,31^. Modelled distributions were then compared to observed frequencies using the Kolmogorov-Smirnov test.

### Genotyping by Sequencing

DNA extracted from lyophilized leaves for each F2 individuals as previously described^72^. DNA concentrations were quantified using the QuantiFluor^®^ dsDNA System kit according to the manufacturer’s instructions (Promega) in the Roche LightCycler 480 (LC480) system. Double digest restriction-site associated DNA libraries were prepared as previously described^73^. A total of 500 ng of high-quality genomic DNA per individual was digested with EcoRI and MspI (New England Biolabs, Ipswich, MA). 50 ng of each digestion reaction was ligated to barcoded adapters and pooled in 48 samples group. Size selection on each pool was performed via gel extractions using NucleoSpin Gel and PCR Clean-up kit (MACHEREY-NAGEL) targeting fragment sizes comprised between 250 and 500 bp. Distinct pools were combined at equimolar ratios and sequenced using the Illumina NextSeq 500 instruments (2 x 75 cycles). Reads were demultiplexed by Illumina index and ddRAD barcode. They were then mapped at per sample level against the *C. rubella* reference genome (Cru_183 from phytozome.org) using BWA-MEM^74^. Variant calling was performed using SAMtools^75^. Those calls were, then, further processed using R^57^ to code the genotypes relative to their parents and select variable sites.

### QTL mapping

Linkage maps were constructed using R/ASMap^76^ with the Kosambi function. Individuals with more than 80% of missing genotype information were excluded and individuals with more than 90% identical genotypes were fused. P-values were adjusted until most of the markers were assigned to a linkage group. In total 1420 and 379 markers were used to reconstitute the *Co* x *Cg* and *Co* x *Cr* genetic map, respectively. Quantitative trait loci (QTL) influencing reproductive and vegetative traits were mapped using R/QTL^71^. The Shapiro-Wilk and Kolmogorov-Smirnov normality tests were used to determine whether the frequency distribution of the data analysed followed a normal distribution (**Table S4**). In *Co* x *Cg* petal length, width, leaf area and pollen number were not normal and transformed to a normal distribution using the Box-Cox transformation. In *Co* x *Cr* petal length, ovule number and pollen number were not normal. Only petal length could be transformed to a normal distribution using the Box-Cox transformation. The transformed data were then used in the QTL analysis. For scored traits such as self-incompatibility, we used QTL mapping function with phenotype model adapted to binary traits. An initial single-QTL genome scan was conducted using Haley-Knott regression to calculate the LOD scores. A genome-wide permutation (n= 1,000) test was, then, conducted to assess the LOD significance threshold (5%). Interactions between QTL were determined using the scantwo function with an Haley-Knott regression and significance thresholds (5%) were tested as above, using genome-wide permutation test (n= 1,000). Significant QTL and their interactions were then used to define QTL model that were assessed using the fitqtl function and the drop-one-term analysis. A Multiple-QTL analysis was then used to refine the QTL model until the best fit was identified (**Table S2**). For all significant QTLs and their interactions estimates of the additive effect and dominance deviation were obtained using the fitqtl function. A 1.5-LOD support interval was used to estimate the position of each QTL.

### Statistical analysis

The phenotypic distributions for each genotype or species are presented with box plots. In these plots, the middle lines correspond to the median, the first and third quartiles are represented by the lower and upper hinges. The whiskers extend until the maximum or minimum values comprised with the 1.5 interquartile ranges. Values beyond this range were considered as outliers and indicated as a dot.

Statistical analyses were conducted in R^57^. The multiple comparisons of phenotypic means were assessed by using Tukey’s HSD post hoc test using the agricolae package add-ons implemented in R^57^. For two-sample comparisons, we used a two-tailed Student’s t-test assuming unequal variances. The null hypothesis was rejected at *p* < 0.05. For the kinetic analysis of petal development, the data are presented as mean ± sem.

## Acknowledgements

We thank Doreen Mäker and Christiane Schmidt for plant care and members of the Lenhard, Bäurle and Rosa groups for discussion and comments on the manuscript. This work was funded by a Deutsche Forschungsgemeinschaft grant (SI1967/2) and The Swedish Research Council (Vetenskapsrådet Ref: 2018-04214).

## Author Contributions

AS and NW, CK designed the project. NW, MC, LA and AS performed the experiments. CK carried out bioinformatic analyses. BN contributed novel biological materials. All authors analysed data. AS supervised the project. AS wrote the manuscript with input from all authors. All authors discussed and commented on the manuscript.

## Supplementary figures

**Fig. S1.**
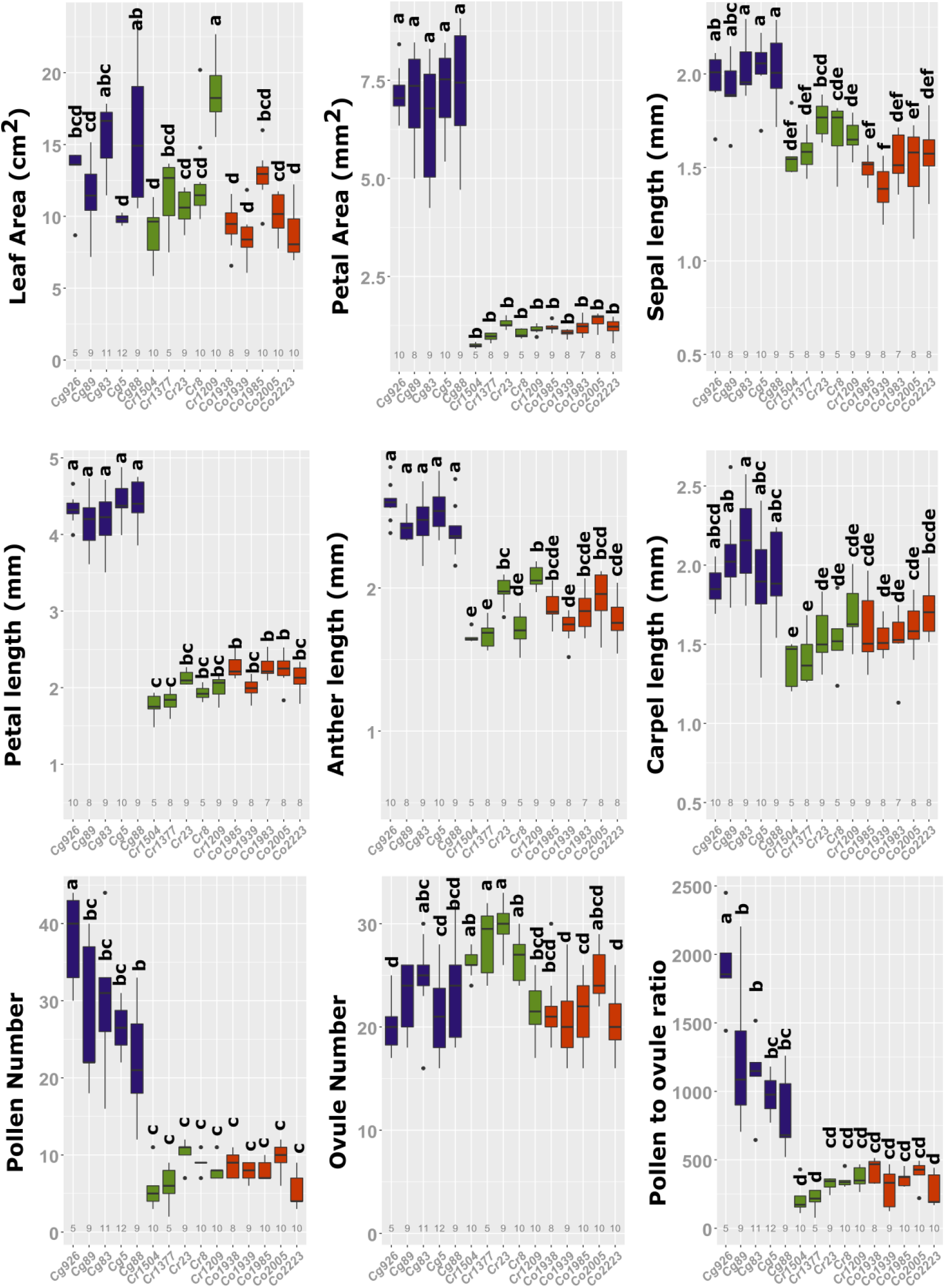
Convergent evolution of the flower morphology after the transition to selfing in the genus *Capsella.* Quantification of organ size and sexual resource allocation in 5 accessions of each of the three *Capsella* species. Letters indicate significant difference as determined by a Tukey’s HSD test. The number of samples used is indicated under each boxplot.

**Fig. S2.**
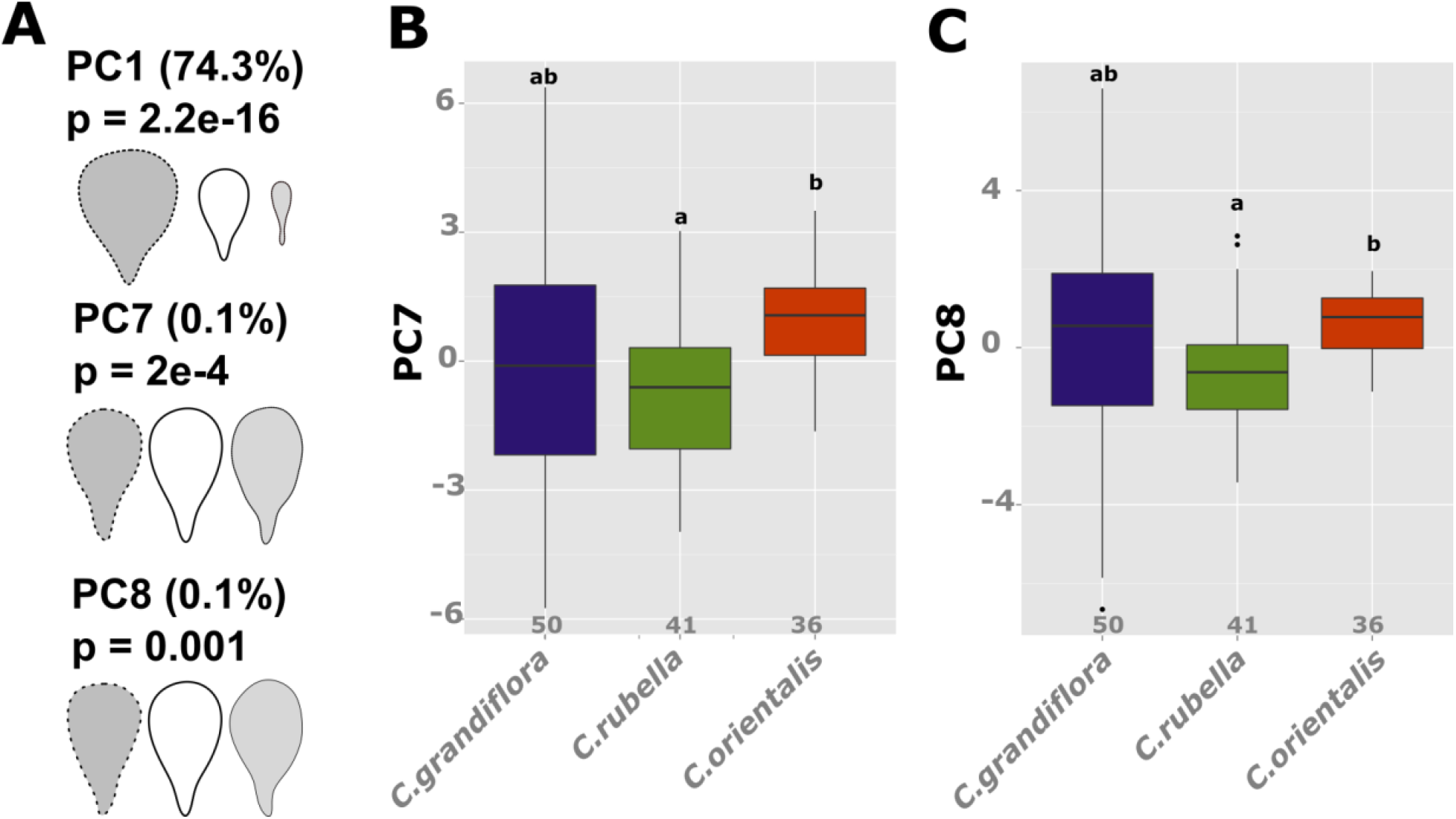
Similar developmental mechanisms underlie petal size reduction in both *C. rubella* and *C. orientalis.* **(A)** The effects along the PC1, 7 and 8 is presented. The percentage of variance explained as well as the *p-value* determined using a Kruskal-Wallis test with the formula PC score ~ species are displayed on the top. **(B)** and **(C)** Distribution of the PC7 (B) and 8 (C) for each genotype. Letters indicate significance as determined by a Tukey’s HSD test. The numbers of samples are indicated under each boxplot.

**Fig. S3.**
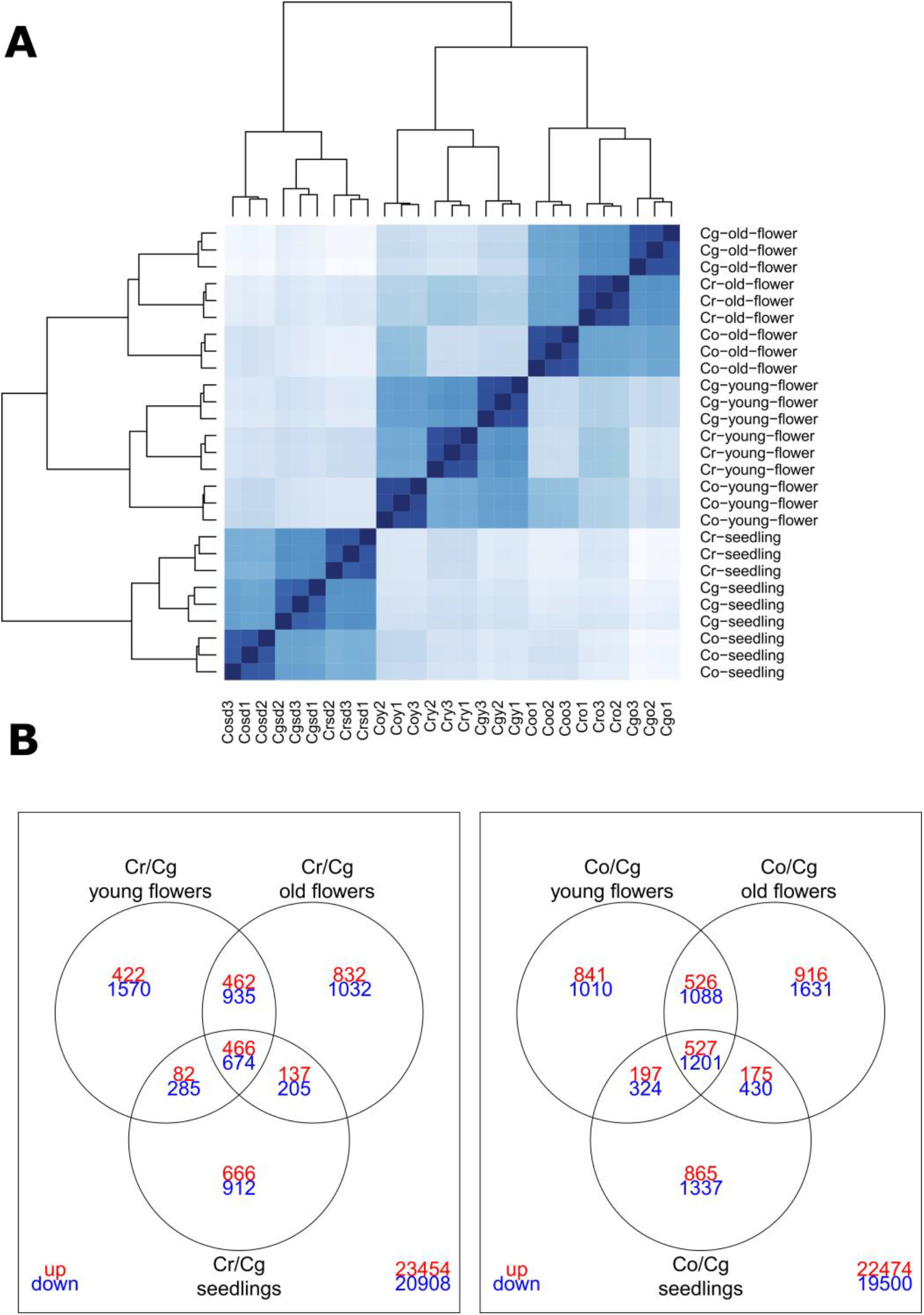
Convergent evolution of gene expression after the transition to selfing in the genus *Capsella.* **(A)** Hierarchical clustering of read counts normalized using a variant stabilization transformation. Note that the samples cluster by replicates, species and tissue-type. **(B)** The list of genes called differentially expressed between *Cr* (panel on the left) or *Co* (panel on the right) and *Cg* are compared between different tissues. The numbers of genes up-regulated are marked in red and the genes down-regulated are indicated in blue.

**Fig. S4.**
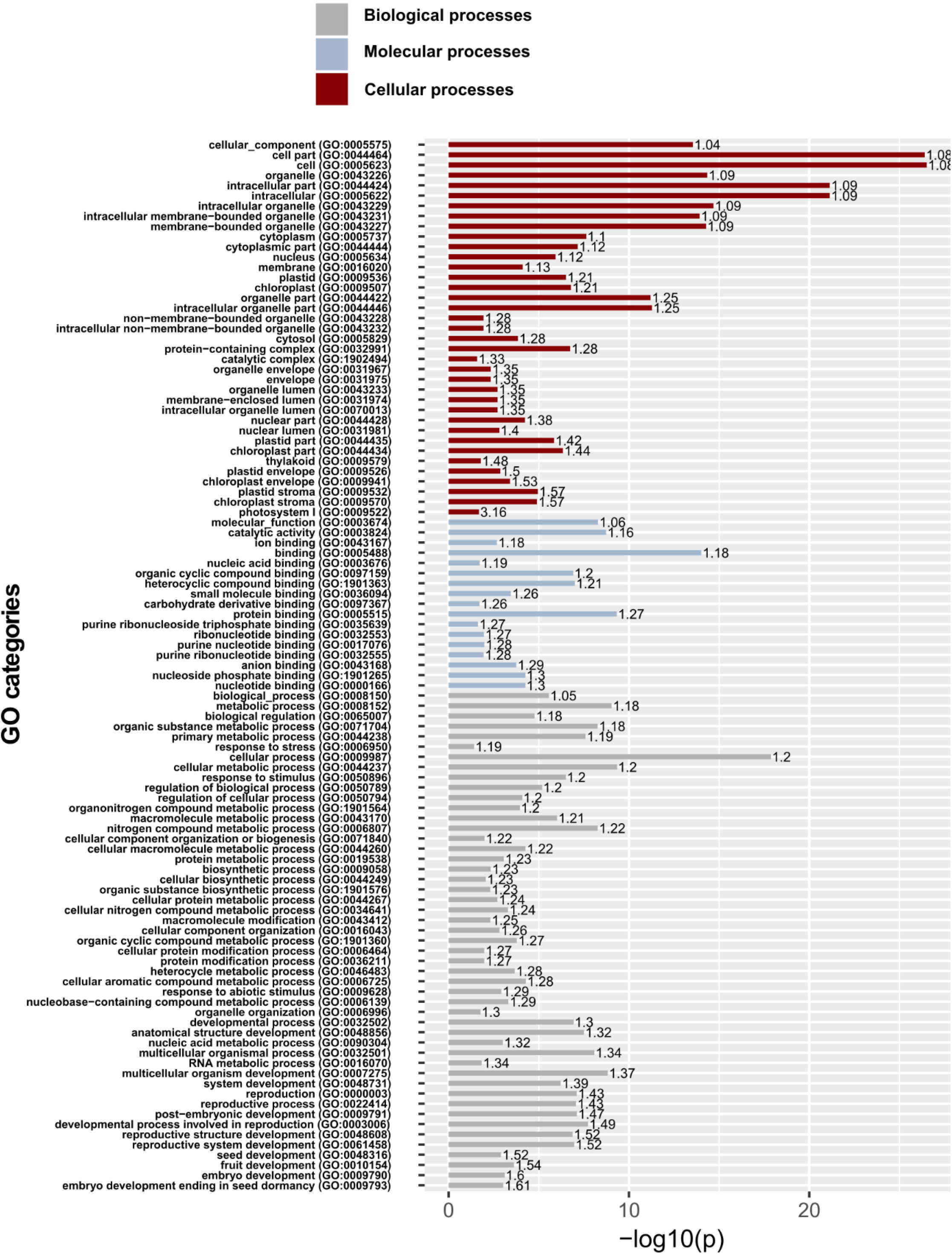
Convergent evolution of gene expression after the transition to selfing in the genus *Capsella.* Gene Ontology enrichment analysis. The gene categories enriched within the genes called differentially expressed in both selfers (coDEGs) are shown for each functional process. The significance (-log10 Fisher’s exact test p-value) of the enrichment is plotted in a bar plot. The number on the top of each bar indicates the fold-enrichment.

**Fig. S5.**
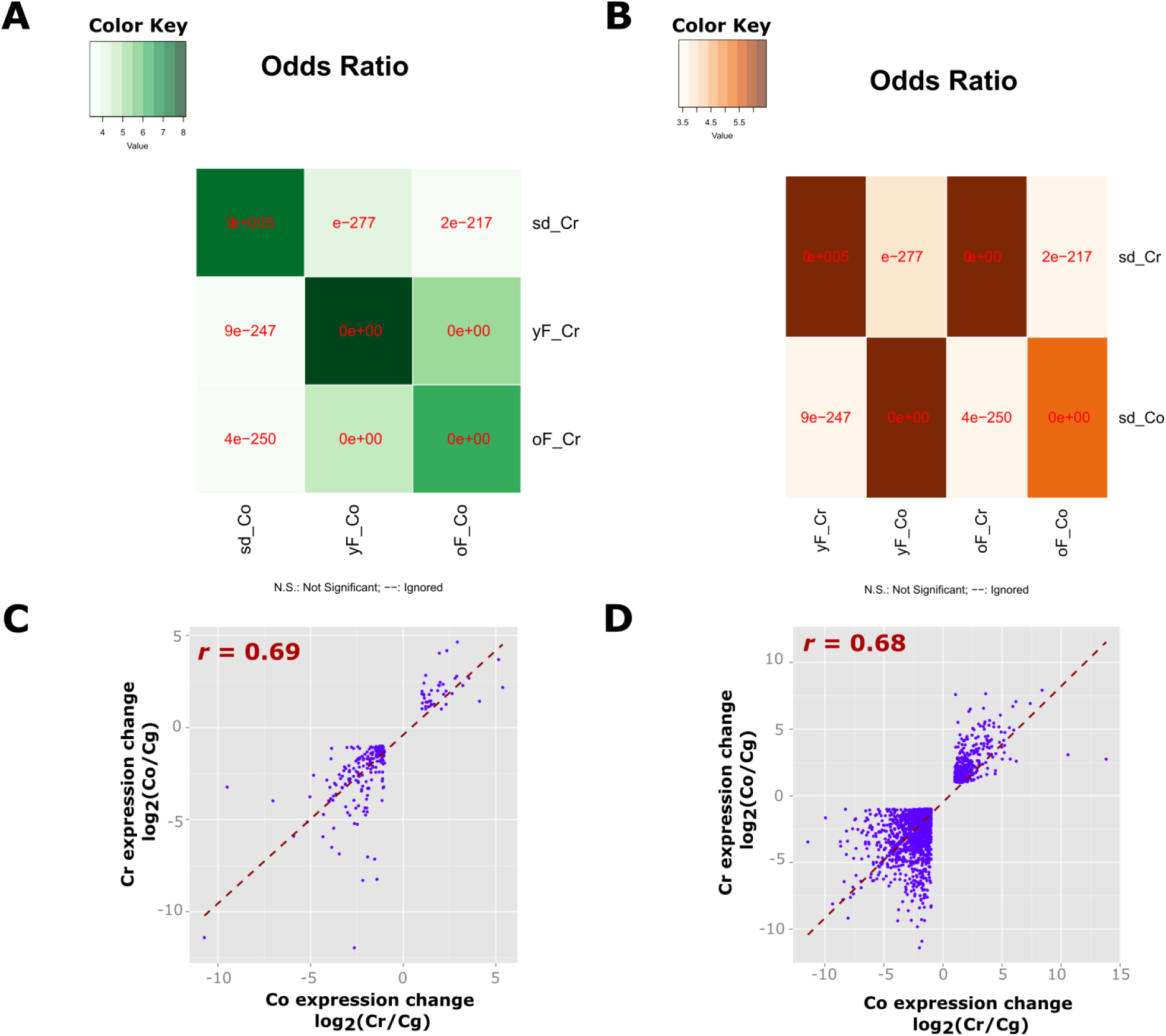
Higher divergence in the flower transcriptome after the transition to selfing. **(A)** and **(B)** Heatmap of Fisher’s exact test odds ratio illustrating the strength of association between the list of genes called differentially expressed between *Cr* and *Cg* and those called differentially expressed between *Co* and *Cg* in all samples (A) or in flowers against seedling samples (B). **(C)** and **(D)** Correlation in gene expression changes between the *Cr* and *Co* DEGs in seedling (C) and flower (D). The Spearman correlation coefficients (r) are indicated on the graphs.

**Fig. S6.**
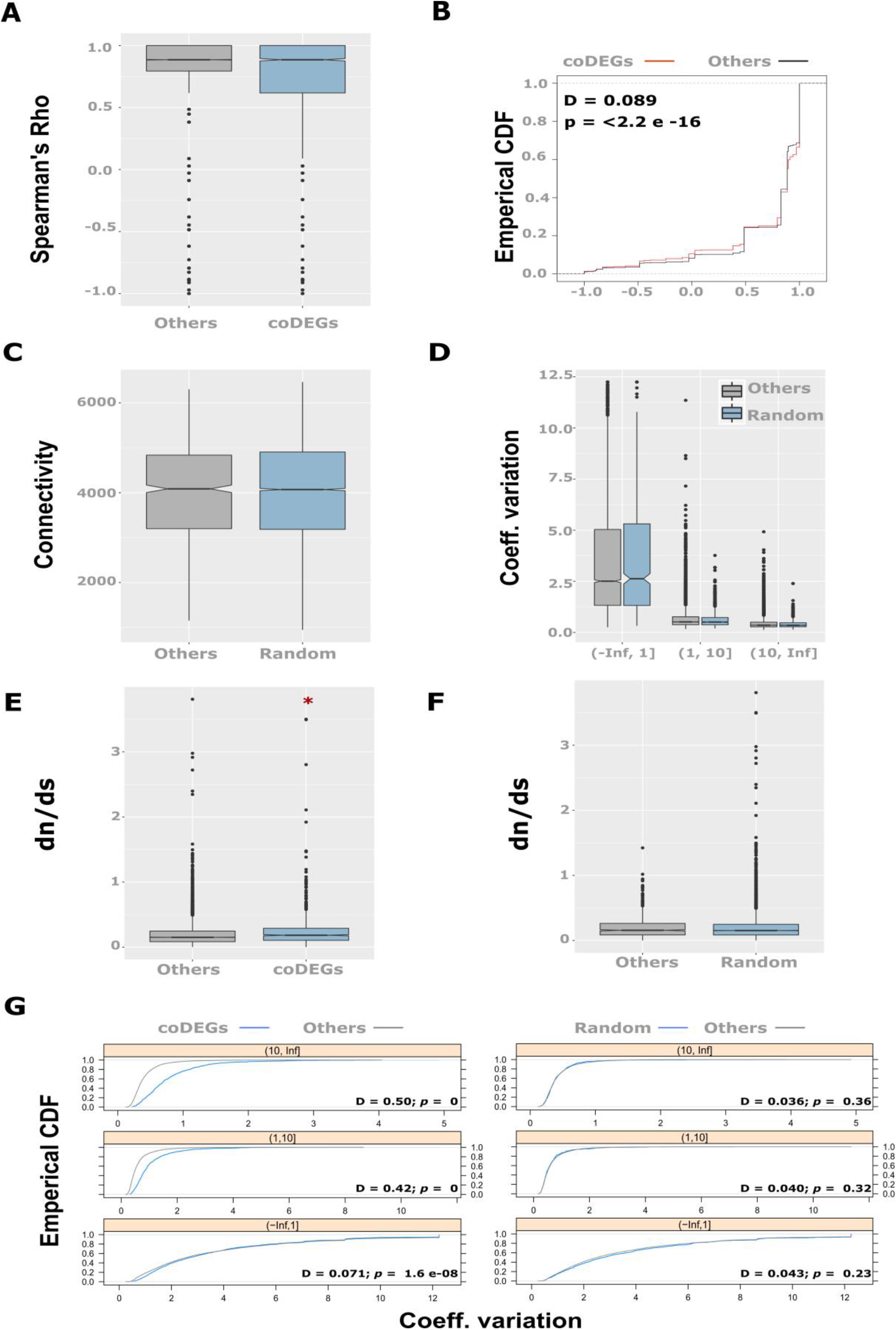
Low-pleiotropic gene regulatory networks underlie the convergent evolution of morphology. **(A)** The distribution of the Spearman’s correlation coefficient of gene expression changes between tissues across *Capsella* species. The list of genes called differentially expressed in the two selfers is compared to those of genes differentially expressed in only one selfer (Others). **(B)** Cumulative distribution functions of spearman’s Rho coefficient of the correlation of gene expression changes between tissues across *Capsella* species. The genes called differentially expressed in the two selfers are compared to genes differentially expressed in only one selfer. The Kolmogorov-Smirnov test was used to compare the two distributions. The returned D-statistic and p-value are indicated on the graph. **(C)** The distribution of the sum of network connectivity (as determined by^36^) for 2000 randomly drawn genes is compared to all other genes (Others). Significant differences between the two distributions were tested using a Wilcoxon test. **(D)** The distribution of the coefficient of variation in expression level within a *Cg* population for 2000 randomly drawn genes (blue) is compared to all other genes (grey). The distributions are determined displayed for each expression class defined based on the mean normalized read count (tags per millions). Significant differences between the two distributions were tested using a Wilcoxon test. **(E)** The distribution of dn/ds ratio (as determined by^36^) for the list of genes called differentially expressed in the two selfers is compared to all other genes. Asterisks indicate significance (p < 0.05) determined by a Wilcoxon test. **(F)** The distribution of dn/ds ratio (as determined by^36^) for 2000 randomly drawn genes is compared to all other genes (Others). Significant differences between the two distributions were tested using a Wilcoxon test. **(G)** Cumulative distribution function of the coefficient of variation of gene expression within *Cg* for coDEGs and all other genes are shown on the left for each expression class. Cumulative distribution function of the coefficient of variation of gene expression within *Cg* for 2000 randomly drawn genes and all other genes are shown on the right for each expression class. The Kolmogorov-Smirnov test was used to compare distributions. The returned D-statistic and p-value are indicated on the graph.

**Fig. S7.**
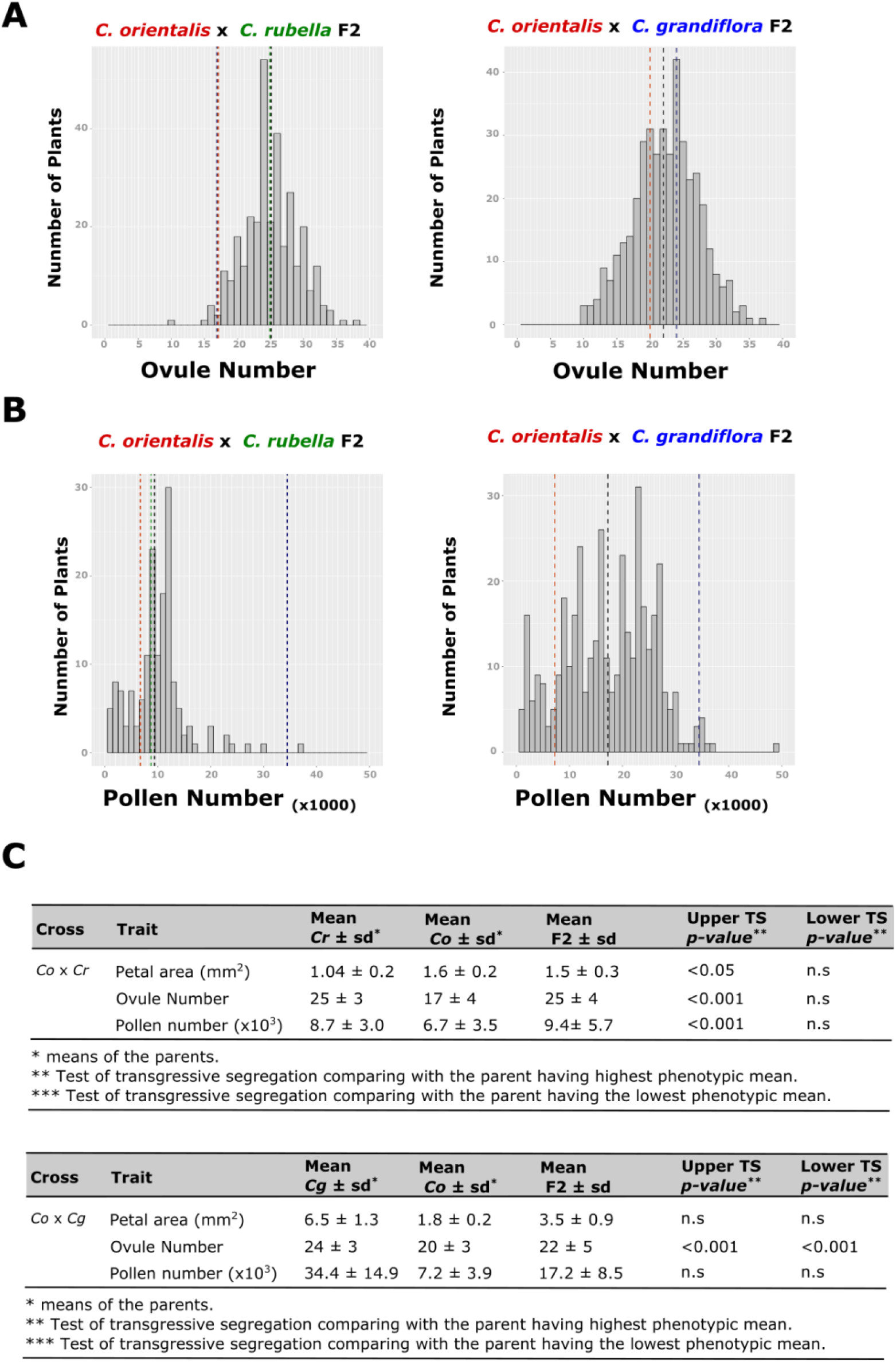
Segregation analysis of selfing syndrome traits. **(A)** Ovule number frequency distribution in *Co* x *Cr* and *Co* x *Cg* F2 population. **(B)** Pollen number frequency distribution in *Co* x *Cr* and *Co* x *Cg* F2 population. **(C)** Summary statistics and transgressive segregation analysis of selfing syndrome traits in *Co* x *Cr* and *Co* x *Cg* F2 population.

**Fig. S8.**
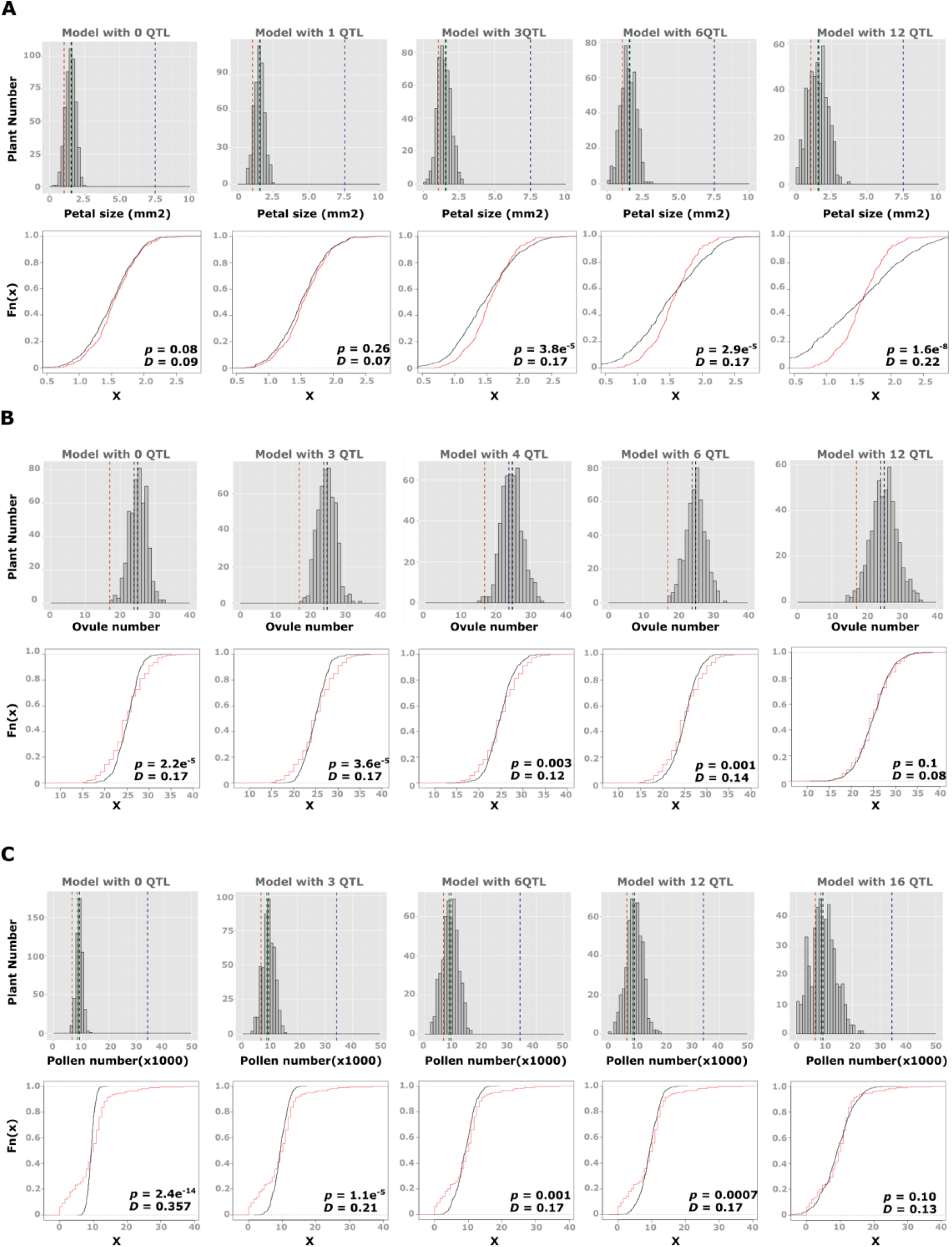
Segregation analysis of selfing syndrome traits. Comparison between observed and simulated petal size **(A)**, ovule number **(B)** and pollen number **(C)** distributions. Modelled distributions are shown on the upper panels. The number of QTL used in the QTL model is indicated on the top. Cumulative distribution function of simulated distributions (in black) is compared to the observed distributions (in red) on the lower panels. The Kolmogorov-Smirnov test was used to compare the two distributions. The returned D-statistic and p-value are indicated on the graph.

**Fig. S9.**
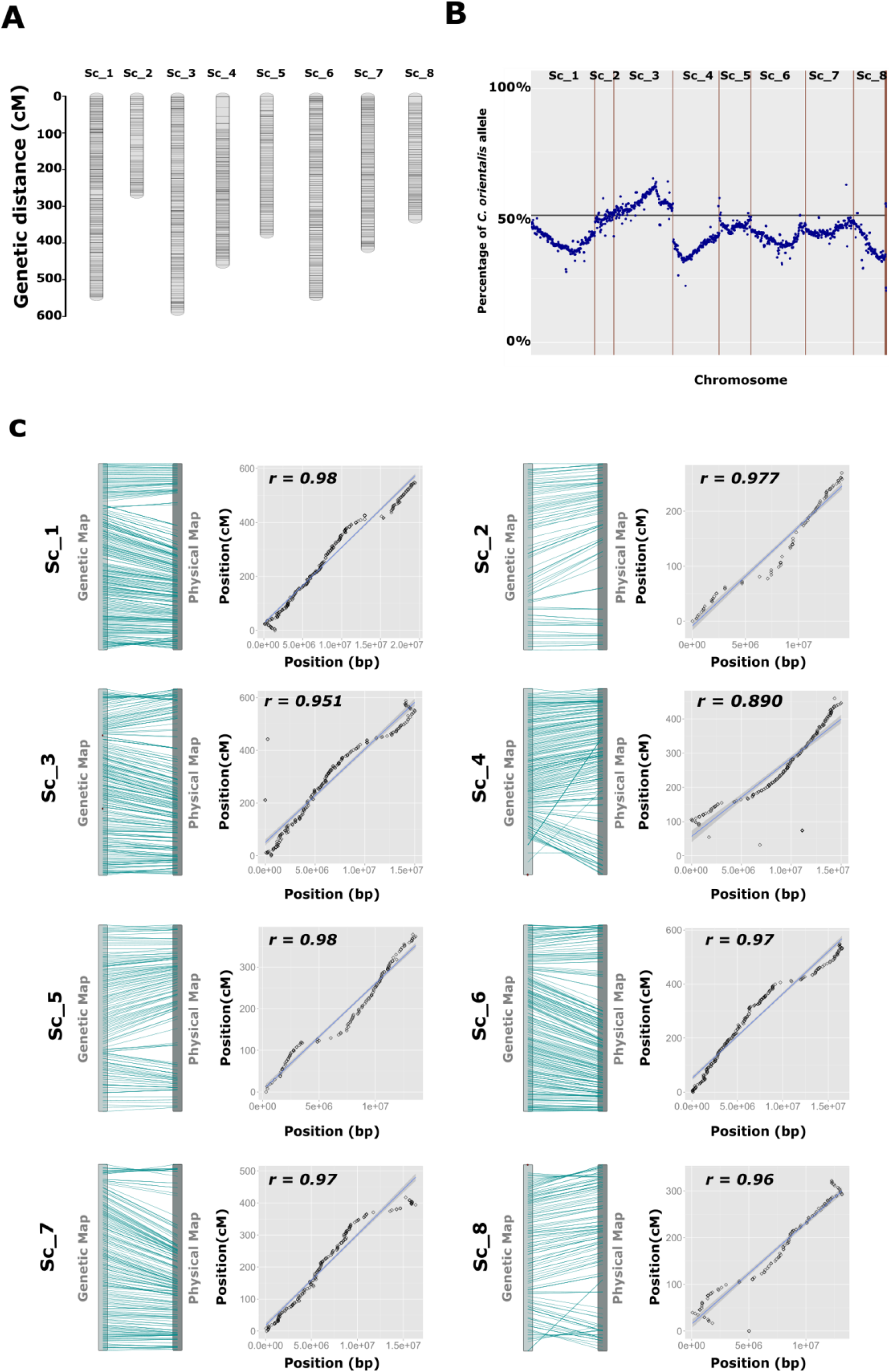
Genetic map of the *Co* x *Cg* population. **(A)** Individual linkage groups are shown in grey bars. Black lines indicate the position of the ddRAD markers. The genetic distances are shown in centimorgan (cM) on the left. **(B)** Segregation distortion. The percentage of *Co* allele estimated at each marker position is shown. **(C)** Comparison of the genetic and physical map. A schematic representation comparing the position of each marker (blue lines) on the physical and genetic map is displayed on the left. A graphical representation of the distance between each marker is shown on the right. The similarity between the two maps was assessed using a Pearson’s correlation test. The returned r coefficients are displayed on the graphs.

**Fig. S10.**
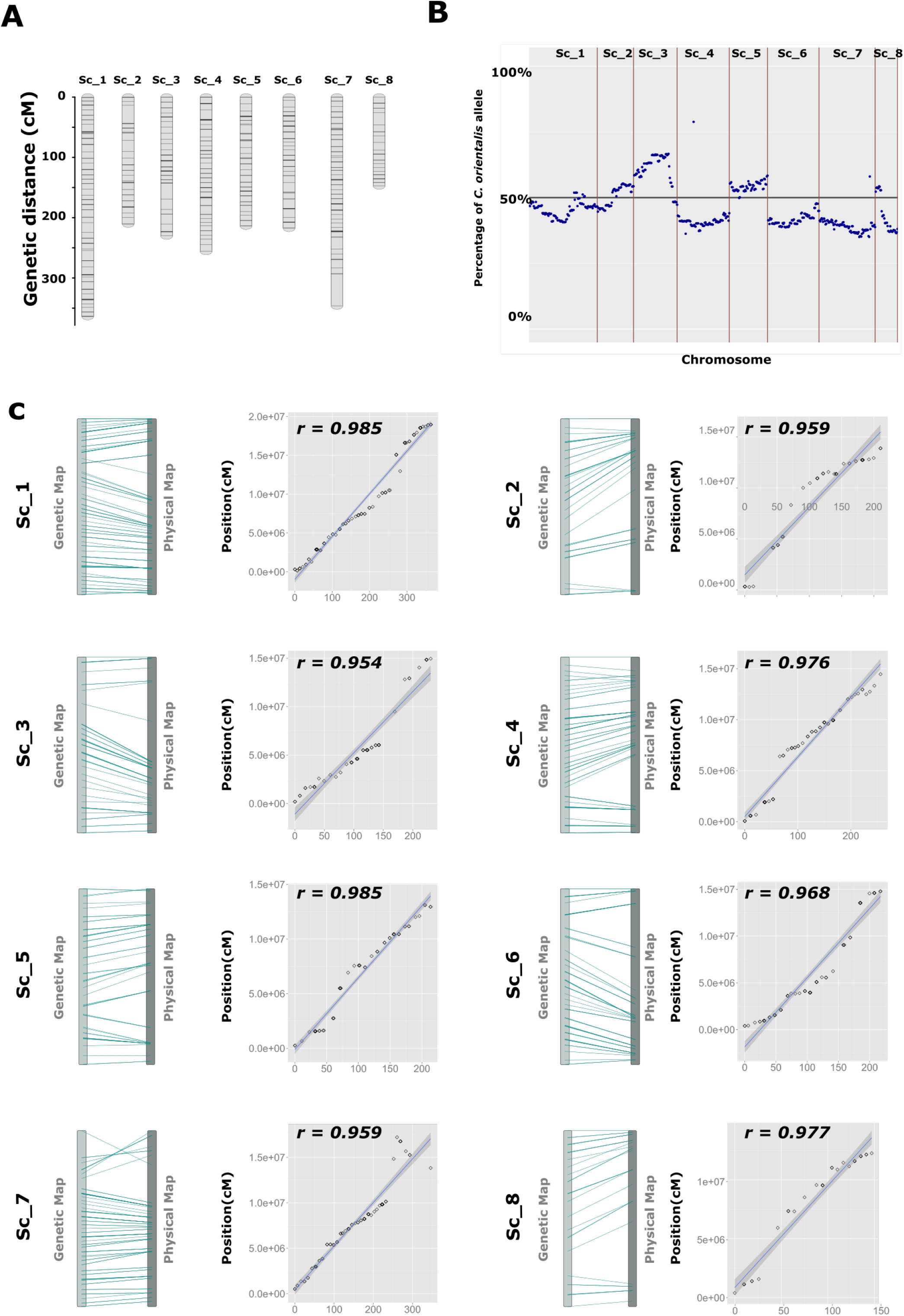
Genetic map of the *Co* x *Cr* population. **(A)** Individual linkage groups are shown in grey bars. Black lines indicate the position of the ddRAD markers. The genetic distances are shown in centimorgan (cM) on the left. **(B)** Segregation distortion. The percentage of *Co* allele estimated at each marker position is shown. **(C)** Comparison of the genetic and physical map. A schematic representation comparing the position of each marker (blue lines) on the physical and genetic map is displayed on the left. A graphical representation of the distance between each marker is shown on the right. The similarity between the two maps was assessed using a Pearson’s correlation test. The returned r coefficients are displayed on the graphs.

**Fig. S11.**
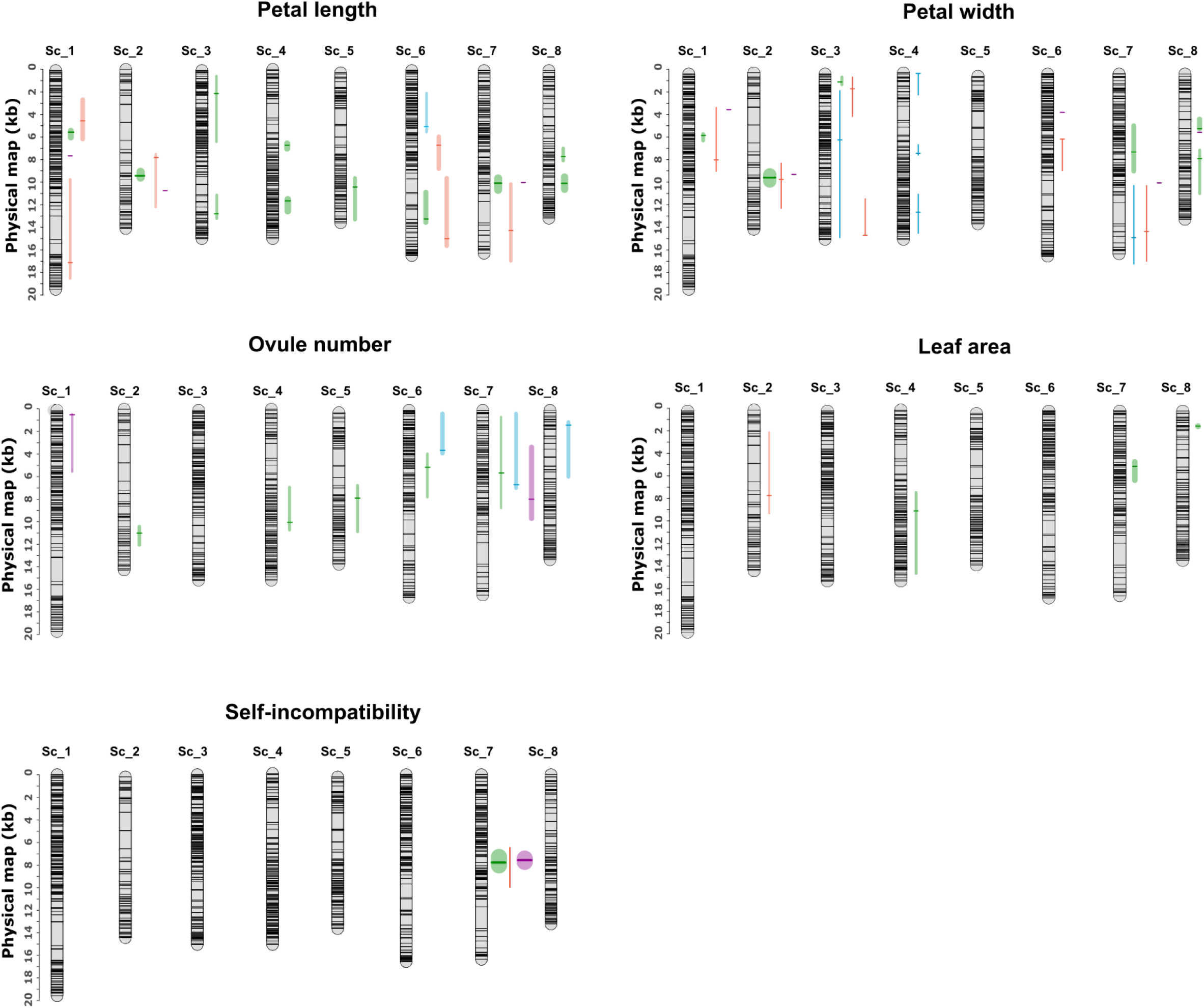
Genetic basis of selfing syndrome traits in *Capsella.* Comparison of the genetic bases of leaf size, petal length and width as well as self-incompatibility between *Cr* and Co. The positions of QTL influencing petal size are indicated on the physical map. The QTL identified in this study in the *Co* x *Cg* population are indicated in green, in the *Co* x *Cr* are indicated in blue. Those previously identified in the *Cr* x *Cg* RILs or F2 populations^25,31^ are shown in red and purple, respectively. The LOD scores peak are indicated by horizontal lines and the vertical lines represent the 1.5 LOD score confident interval. The width of the vertical lines illustrates the strength of the QTL.

## Supplementary tables

**Table S1.** Geographic origin of accessions used in this study

| Species | Designation | Country of origin, locality | Coordinates | Provided |
| --- | --- | --- | --- | --- |
| *C. grandiflora* | *Cg926* | Greece; Votonosi | 39°46N;<br>21°70E | B. Neuffer |
|  | *Cg89* | Greece; near Koukouli, Zagori | 39°52N;<br>20°46E | T. Slotte |
|  | *Cg83* | Greece; near Vathyrevma | 39°26N;<br>21°25E | T. Slotte |
|  | *Cg5* | Greece, near Troumpeta, Corfu | 39°42N;<br>19°45E | T. Slotte |
|  | *Cg88* | Greece; Monodendri, Zagori | 39°53N;<br>20°44E | T. Slotte |
| *C. rubella* | *Cr1504* | Spain; Canary Islands; La Palma | 28°40N;<br>17°52W | B. Neuffer |
|  | *Cr1377* | Argentina; Province of Buenos Aires | 37°20S;<br>59°84W | T. Slotte |
|  | *Cr23* | Italy; Senerchia | 40°70N;<br>15°20E | T. Slotte |
|  | *Cr8* | Luxembourg; Esch-sur-Sûre | 49°90N;<br>05°93E | T. Slotte |
|  | *Cr1209* | Spain; Cumbre Dorsal, Tenerife | 28°29N;<br>16°60W | T. Slotte |
|  | *Cr4.23* | Pola de Somiedo; Spain | 43°092N;<br>06°25W | T. Slotte |
| *C. orientalis* | *Co1938* | Russia; Siberia, Altai Kraj, city of Barnaul | 53°20N;<br>83°45E | B. Neuffer |
|  | *Co1939* | Kazakhstan; Pavlodarskaya Oblast, Pavlodar, 400 km north-east from 400km north-north-east from Astana, ruderal in lawn | 52°16N;<br>76°57E | B. Neuffer |
|  | *Co1983* | Russia; Siberia, Altai Kraj; Smeinogorskraion, Kolyvanskoe Lake | 51°22N;<br>82°12E | B. Neuffer |
|  | *Co1985* | Russia; Siberia. Altai Kraj; city of Barnaul | 53°21N;<br>83°44E | B. Neuffer |
|  | *Co2005* | China; Xinjiang, Dzungaria, 485 km north of Urmuchi, Mongolian | 48°05N;<br>88°56E | B. Neuffer |
|  | *Co2223* | unknown | unknown | B. Neuffer |

**Table S2.** Multiple-QTL models

| Population | Trait | Model | LOD | % Variance explained |
| --- | --- | --- | --- | --- |
| *Co1983 x Cg926* F2 | **Petal area** | PAQTL1 + PAQTL2 + PAQTL3 + PAQTL4 + PAQTL5 + PAQTL6 + PAQTL7 + PAQTL8 + PAQTL5:PAQTL8 | 52.06 | 53.06 |
|  | **Petal length (boxcox)** | PLQTL1 + PLQTL2 + PLQTL3 + PLQTL4 + PLQTL5 + PLQTL6 + PLQTL7 + PLQTL8 + PLQTL9 + PLQTL10 + PLQTL11 + PLQTL1:PLQTL8 + PLQTL9:PLQTL11 + PLQTL5:PLQTL6 | 64.17 | 60.52 |
|  | **Petal width (boxcox)** | PWQTL1 + PWQTL2 + PWQTL3 + PWQTL4 + PWQTL5 + PWQTL6 | 44.19 | 47.27 |
|  | **Ovule number** | OQ1 + OQ2 + OQ3 + OQ4 + OQ5 | 21.57 | 22.14 |
|  | **Leaf area (boxcox)** | LAQ1 + LAQ2 + LAQ3 + LAQ2:LAQ3 | 17.44 | 17.91 |
| *Co1983 x Cr4.23* F2 | **Petal area** | PAQTL1 + PAQTL2 | 7.97 | 11.6 |
|  | **Petal width** | PWQTL1 + PWQTL2 + PWQTL3 + PWQTL4 + PWQTL5 | 18.66 | 25.2 |
|  | **Ovule number** | OQ1 + OQ2 + OQ3 + OQ1:OQ3 | 16.5 | 21.87 |
| Epistatic effects between QTL are presented as “:.” |  |  |  |  |

**Table S3:**
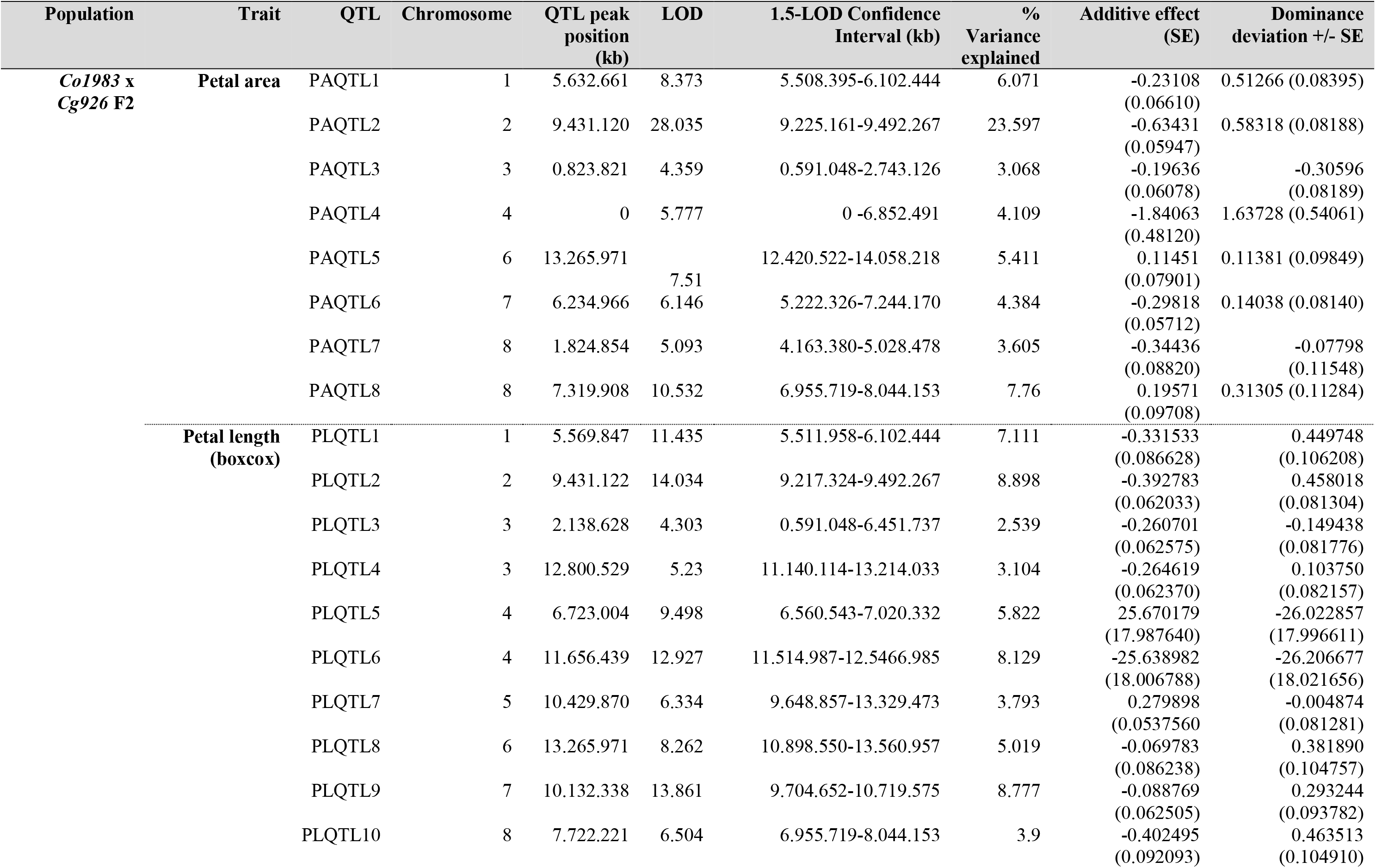
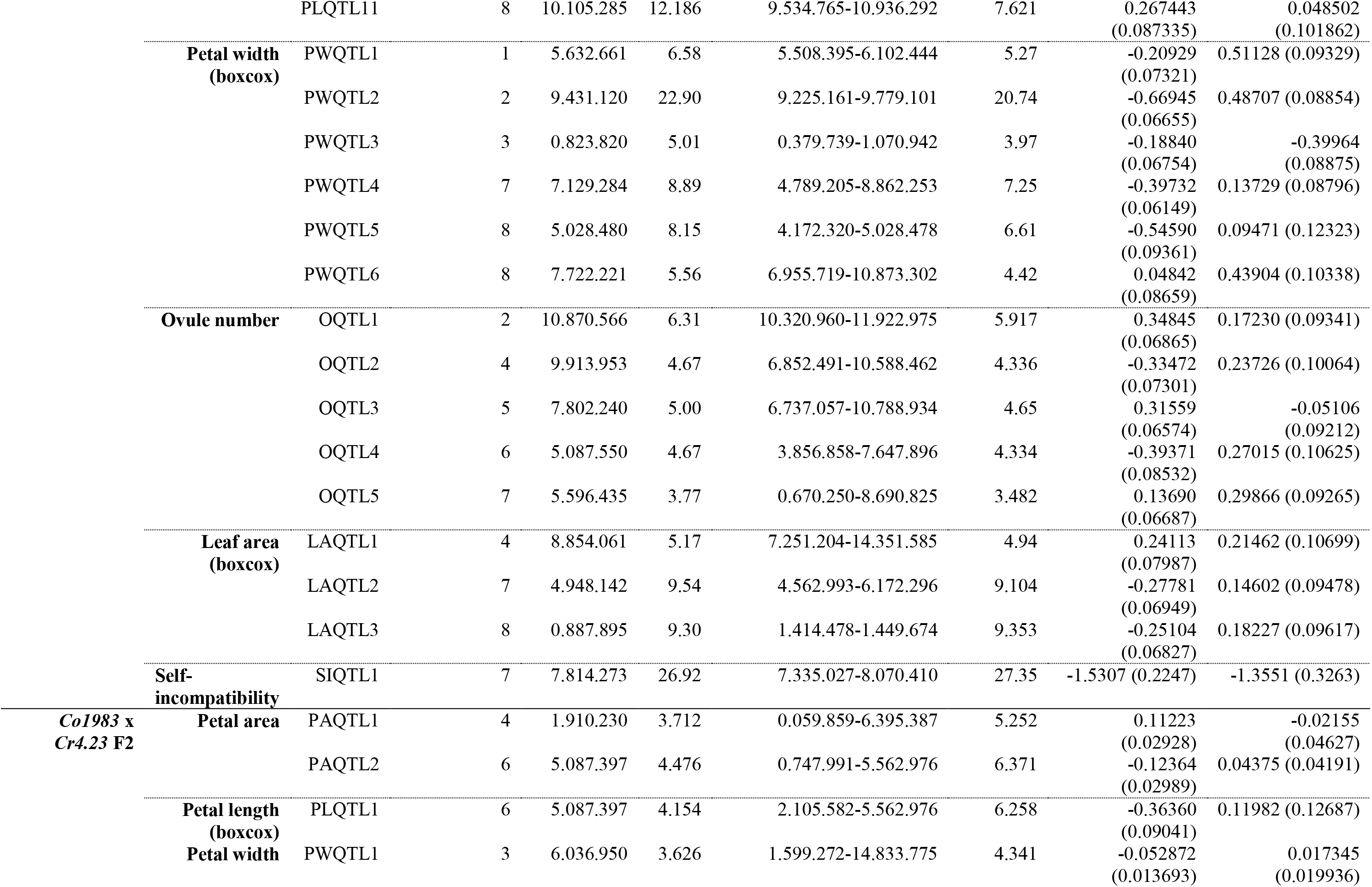
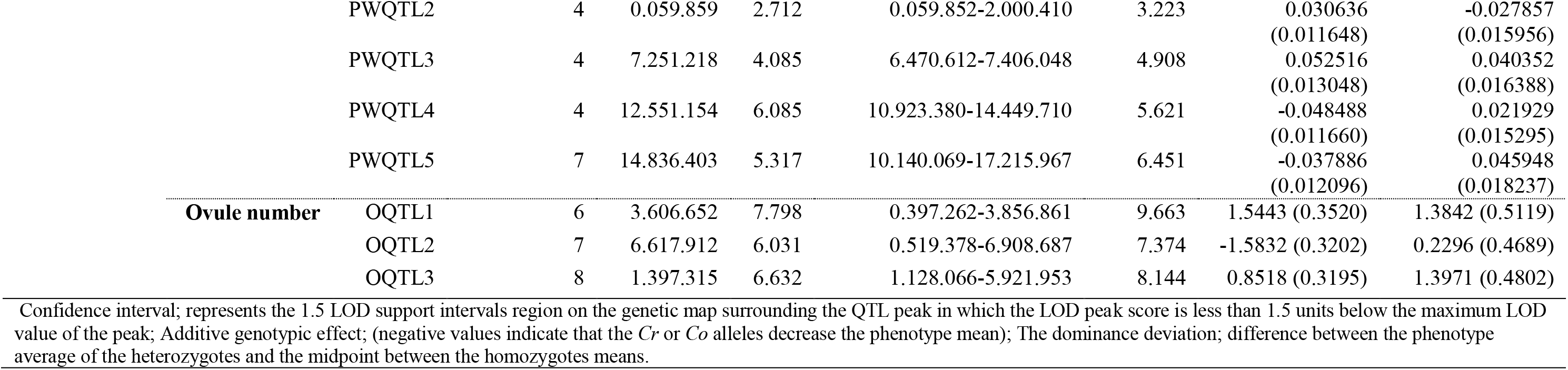
QTL analysis

**Table S4.**
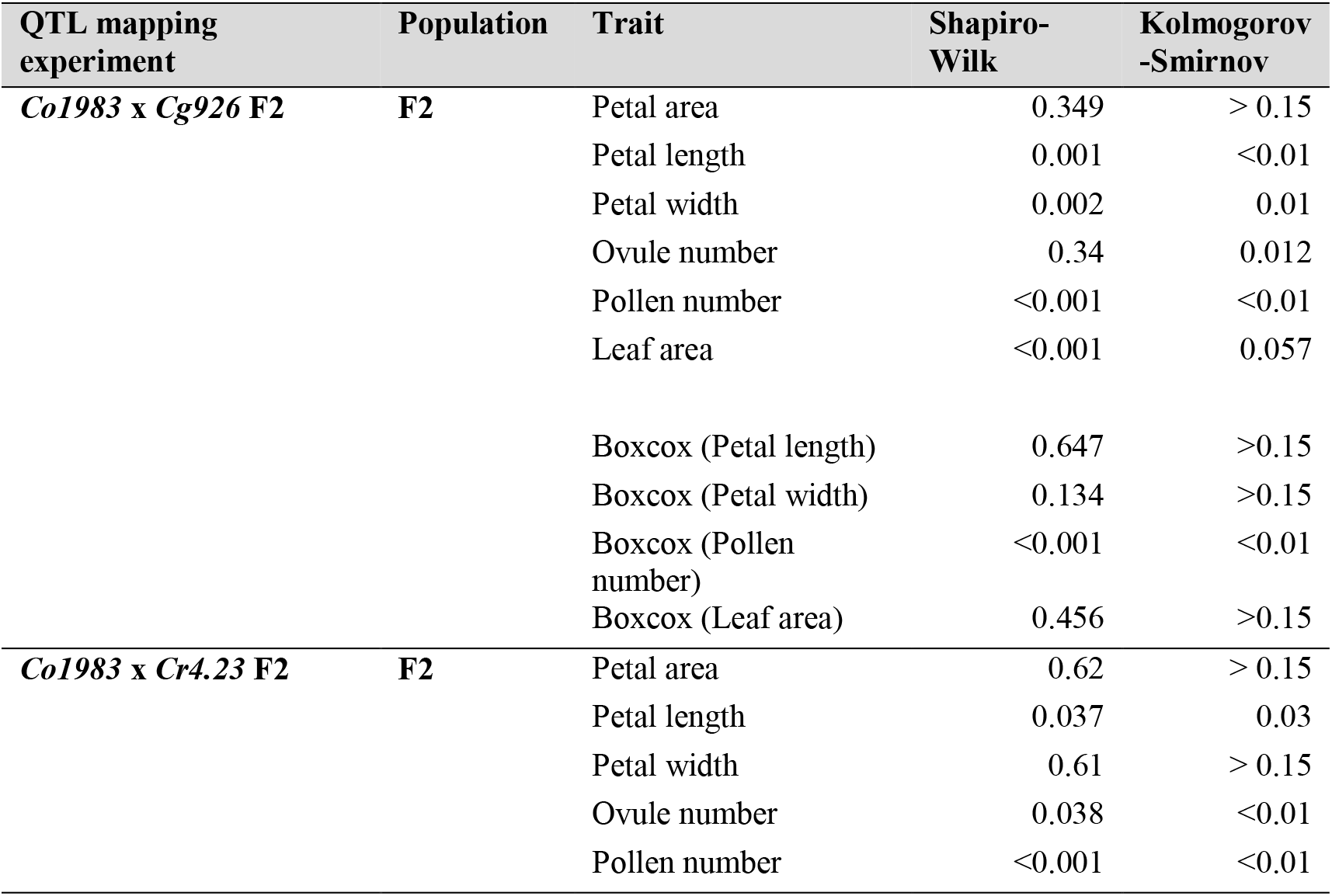
Normality tests for distribution of phenotypes in the F2 and parental populations

| QTL mapping experiment | Population | Trait | Shapiro-Wilk | Kolmogorov-Smirnov |
| --- | --- | --- | --- | --- |
| *Co1983* x *Cg926* F2 | F2 | Petal area | 0.349 | > 0.15 |
|  |  | Petal length | 0.001 | <0.01 |
|  |  | Petal width | 0.002 | 0.01 |
|  |  | Ovule number | 0.34 | 0.012 |
|  |  | Pollen number | <0.001 | <0.01 |
|  |  | Leaf area | <0.001 | 0.057 |
|  |  | Boxcox (Petal length) | 0.647 | >0.15 |
|  |  | Boxcox (Petal width) | 0.134 | >0.15 |
|  |  | Boxcox (Pollen number) | <0.001 | <0.01 |
|  |  | Boxcox (Leaf area) | 0.456 | >0.15 |
| *Co1983* x *Cr4.23* F2 | F2 | Petal area | 0.62 | > 0.15 |
|  |  | Petal length | 0.037 | 0.03 |
|  |  | Petal width | 0.61 | > 0.15 |
|  |  | Ovule number | 0.038 | <0.01 |
|  |  | Pollen number | <0.001 | <0.01 |

